# *In situ* structure of a gap junction – stomatin complex

**DOI:** 10.1101/2025.03.27.645584

**Authors:** Nils Rosenkranz, Konstantin Wieland, Alexandra N. Birtasu, Sina Manger, Abhishek Bhattacharya, Achilleas S. Frangakis, Alexander Gottschalk

## Abstract

Gap junctions (GJ) are intercellular channels that mediate electrical signals and the transfer of small molecules. GJs are crucial for the functions of the brain, heart and other organs. While structures of purified homomeric GJs are available, we lack *in situ* structures. *In vivo*, GJs can form heteromers with different functionalities, and may associate with other proteins. Here, we analyzed *Caenorhabditis elegans* GJs by cryo-electron tomography and sub-tomogram averaging. We observed hexagonal arrays of GJs at cellular junctions in primary embryonal cell culture that displayed distinct wide and narrow conformations. Moreover, in about 20% of the observed channels, we found a cap-like, cytosolic protein assembly enclosing the channel pore. We propose that the cap-structure is formed by the stomatin UNC-1, which is known to interact with *C. elegans* GJs, and strengthen this hypothesis by matching AlphaFold3 models of UNC-1 multimers with our GJ average. Furthermore, expressing UNC-1 and the *C. elegans* innexin UNC-9 in HEK cells resulted in similar structures at cell-cell contacts. UNC-1/stomatin ring assemblies may affect GJ formation or functions like rectification, that might be evolutionarily conserved.

**Significance Statement:** Gap junction (GJ) channels connect neighboring cells. Structures of (purified) GJs have been studied *in vitro*, but not *in situ*. We identified GJ channels in primary *Caenorhabditis elegans* cells by cryo-electron tomography, and analyzed their structure by sub-tomogram averaging. The channels transverse the membranes of connected cells, and AlphaFold3 (AF3) models of the GJ subunit UNC-9, assuming dodecamers, fit the experimentally obtained surface map well. We observed a cytosolic ‘cap’ structure on the GJ channels. The stomatin protein UNC-1 is known to physically interact with UNC-9 GJs. AF3 models of UNC-1 hexadecamers fit the cap structure, indicating that it may be formed by UNC-1, providing a first idea how UNC-1 interacts with, and may functionally influence, GJ channels.

## Introduction

Multicellular organisms crucially rely on intercellular connections and communication, facilitated by various types of junctions. Communication between excitable cells is largely executed via two forms of synapses. Chemical synapses require neurotransmitters and neuropeptides to act as messengers between cells. Electrical synapses, also known as gap junctions (GJs), form intercellular channels reaching through a gap of 2-4 nm (1, 2). They can synchronize or coordinate activity in excitable cell ensembles (3-5), and they allow the direct exchange of small molecules (metabolites and signaling molecules) (6-9). GJs ensembles have been observed *in situ* by electron microscopy (10) and by fluorescence microscopy, while their functional properties have been studied by electrophysiology (11, 12). In each of the connected cells, GJ channels are formed by 6 subunits, so-called connexins in vertebrates, or 6-8 subunits called innexins in invertebrates (13-17). In principle, gap junctions can be as small as a single channel, however, they often form larger patches at the interface of two connected cells, thus establishing a substantial junction (18). GJs can be homomeric (all subunits identical), or they form heteromeric (different subunits per connexon), as well as homo- or heterotypic channels (non-identical hemichannels in the two connected cells) (12). GJs are constitutively open at rest, however, when the junctional potential or cytosolic Ca^2+^ concentration increase, they close by blocking their pores with their N-termini (19). Based on their subunit composition, or due to posttranslational modification, GJs exhibit differential gating properties and they can achieve rectification (11, 19). Also hemi-channels exist that function independently of GJ counterparts (20-22). In vertebrates, these consist of a third class of subunits, called pannexins, which share ∼20% sequence homology with the invertebrate innexins (13, 23, 24). Pannexin hemichannels were first described to be hexameric like connexins (25, 26). However, a different study suggested an octameric multimer (27), and recent advances in cryogenic electron tomography (cryo-ET) revealed that pannexins can form heptamers (28-30). Some connexin or innexin homo-oligomers were analyzed by crystallography or cryo-EM of overexpressed, isolated GJ channels in detergent or in nanodiscs (16, 31-33), where evidence for hexamers but also for octameric assemblies was found. It is unclear if these differences may depend on different biochemical preparations and reconstitution conditions. *In situ* structural information of GJs and the complexes they form is lacking.

Apart from the structural subunits, GJ function is further regulated by associated proteins like stomatins (34, 35), and scaffolds that interact with the connexons/innexons (36, 37). Yet, the structural interplay of such proteins with the GJ channels is unknown. Recently, the structure of a GJ analog in cyanobacteria was determined (38). These so-called septal junctions form tubes connecting adjacent cells. The diameter is larger than that of eukaryotic GJs, however, they serve a similar purpose (38-40). At each end, septal junctions possess a cytosolic cap and a plug, which can seal the channel upon stress detection (38).

The nematode *Caenorhabditis elegans* possesses 25 innexin genes, with high structural similarity to vertebrate connexins. Innexins are expressed in virtually all cell types in *C. elegans*, but mostly connecting excitable cells (41, 42). The most abundantly expressed *C. elegans* innexins are UNC-7 and UNC-9. They co-localize extensively along the ventral and dorsal nerve cords and are required for coordinated locomotion (34, 43). Some splice variants of UNC-7 convey rectification in heterotypic GJs with UNC-9, which enables directional signal transmission (11). GJs comprising UNC-9 subunits functionally and physically interact with associated UNC-1 proteins (34, 35), where the latter was shown by bimolecular fluorescence complementation (BiFC) assays. UNC-1 is a protein of the stomatin, prohibitin, flotillin and HflK/C (SPFH) domain family, with two predicted N-terminal hydrophobic / transmembrane domains (44-46). While an N-terminal TM helix has been observed in the biological structure of the bacterial HflK/C complex, flotillins lack such a TM helix and bind to the membrane surface by peripheral attachment through lipid anchors and penetration into the outer membrane leaflet (44, 46). The membrane topology of UNC-1 is unknown, i.e. whether the N-terminal helix traverses the membrane; the second hydrophobic domain resembles the flotillin SPFH1 domain that peripherally penetrates the membrane (**Fig. S10**). *unc-1* mutations can act in a dominant negative manner on UNC-9 GJs (34, 35, 47). UNC-1 may interact also with other GJs, or have functions extending beyond its interaction with UNC-9.

Here, we studied the *in situ* structure of *C. elegans* GJs by cryo-ET and sub tomogram averaging (STA). We observed both narrow and wide (possibly closed and open) channels in hexagonal arrangements. In addition, we find that some GJs carry a cytosolic cap structure, either in one, or in both connected cells. Following the hypothesis that this cap may be constituted by the stomatin protein UNC-1, we used AF3 (48) modeling to analyze the putative structure of UNC-1. The stomatin forms a ring-like, multimeric structure, and in its hexadecameric form, resembles the surface model of the cytosolic cap we observed by STA.

## Results

### Targeting cellular junctions in primary cell culture

To study the *in situ* structure of GJs, we tested different approaches toward identifying and preparing them in *C. elegans*. We used a correlative light and electron microscopy (CLEM) approach, labeling markers for these junctions with fluorescent proteins, and preparing regions where these markers are enriched for (label-free) cryo-ET. Samples for cryo-ET of unstained tissue need to be very thin (<200 nm) to preserve high-quality structural information (49). Thus, we grew embryonal primary cultures from *C. elegans* larvae on carbon coated EM grids, assuming that these cells may form relevant synaptic contacts, including GJs, and identified regions containing junctions by fluorescence microscopy. Neuronal processes (neurites) grown in culture generally are sufficiently thin to enable direct observation by cryo-ET (**Fig. 1A, B**). We used several transgenic *C. elegans* strains expressing fluorescent protein fusions, marking synapses in *C. elegans* neurons (**Fig. S1A**). In embryonal stages, the proportion of GJs relative to chemical synapses is high, making the formation of mixed chemical and electrical synapses (and thus, visible colocalization) more likely (50, 51). We thus used markers of GJs (innexins UNC-9 and UNC-7), and of chemical synapses (the active zone marker ELKS-1 in cholinergic neurons; the synaptic vesicle-associated small G protein RAB-3 in all neurons). In addition, we targeted the stomatin-like protein UNC-1, which functionally and physically interacts with UNC-9 (34, 35). We examined primary embryonal cells of the transgenic animals cultured on EM carbon grids by fluorescence microscopy (**Figs. 1B; S1A, B**). The fluorescently tagged proteins reliably showed the expected punctate expression pattern in the nervous system of adult animals, as well as in the respective embryonal culture, at cell-cell contacts of the thin protrusions representing neurites.

**Figure 1:**
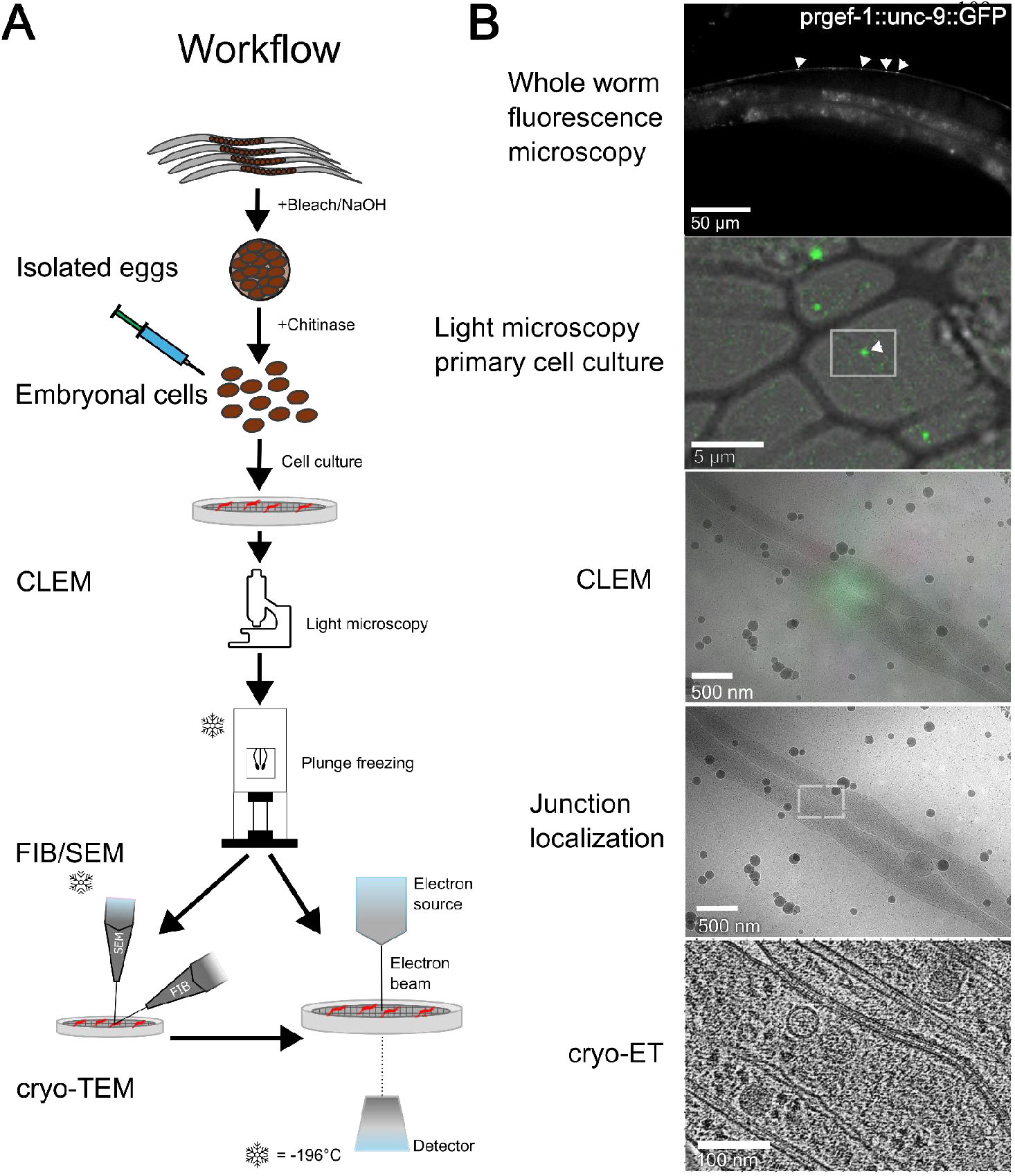
Workflow for preparation, and *in situ* cryo electron tomography, of cellular junctions. **(A)** Embryos from *C. elegans* expressing fluorescently labeled GJ subunits or other marker proteins are dissociated to grow cell cultures on EM grids. Fluorescence microscopy is used to identify regions of interest, cells are plunge-frozen and the regions observed by cryogenic transmission electron tomography (cryo-ET). If the sample is too thick, focused ion beam milling (FIB) can be used to cut thin lamellae. **(B)** Example images of the indicated stages of the process to find cellular junctions by cryo-ET. Bottom image shows a cell-cell junction of very low spacing (<5 nm). Scale bars are indicated in each panel.

Plunge-frozen samples of these fluorescently labelled cell processes were examined with cryo-ET after specific regions were selected by CLEM. On regions of interest that appeared to be too thick for direct visualization by cryo-TEM, an additional thinning step was performed by cryo-focused ion beam (FIB) milling. The cellular proteome including ribosomes, the cytoskeleton with actin filaments and microtubules, and various other complexes could be visualized in the vicinity of the cell-cell junctions (**Figs. 1B; S1B; S2A-F**). Specifically, at the sites of fluorescence, GJs were characterized by membrane distances of ca. 5 nm (**Figs. 1B; 2A, B, D**) and protein densities that appeared to span both membranes, with a cytosolic connection and a channel throughout (**Fig. 2A, B, D)**. We also observed apparent top views of GJ patches containing several channels (**Figs. 2B, C; S3**). Upon closer inspection, in both the side- and the top views, we noticed that about half of the channels were wide, while the others were narrow (**Figs. 2C, D; S3**).

**Figure 2:**
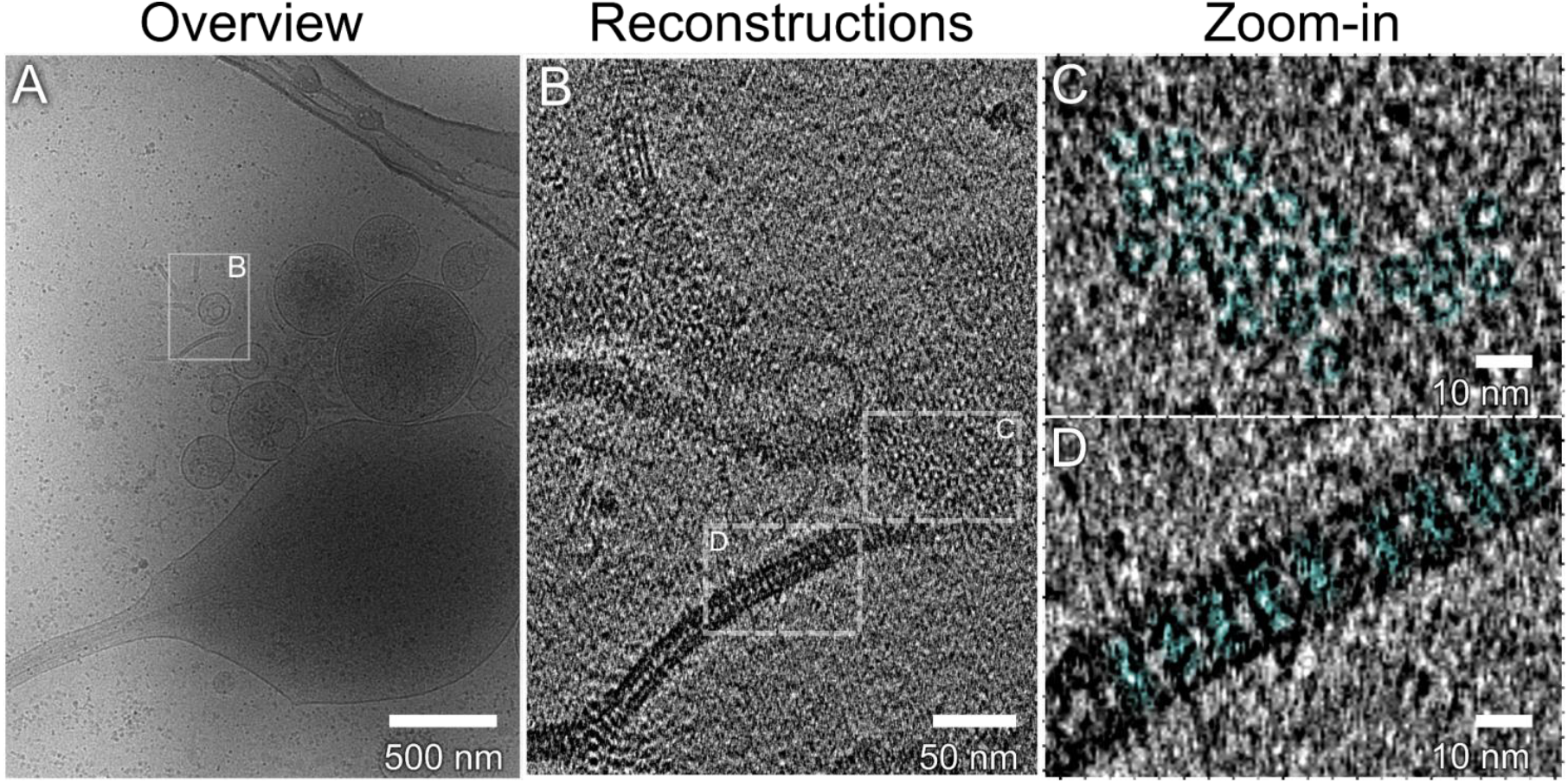
Gap junctions can be identified by cryo-ET of cultured *C. elegans* cells. **(A)** Overview, low magnification cryo-TEM image of a region of an EM grid. Box highlights region containing cell-cell contacts. Note that only membranes perpendicular to the sample plane can be seen. **(B)** Higher magnification cryo-ET images of the box in (A). **(C, D)** Gap junctions are seen in side view (B, D) and top view (B, C). High magnification images, indicated as boxes in (B). Top views indicate channels with a ‘wide’ configuration, see **Fig. S3** for other channels adopting a ‘narrow’ configuration. Individual GJ channels are highlighted in transparent cyan. Scale bars are indicated in each panel.

### Heterogeneity of GJ channels suggests open and closed states and hexagonal arrangement in GJ patches

We next sought to determine the structure of these channels by STA. We manually classified the sub tomograms into wide and narrow channels, yielding a final dataset of 562 wide junctions (272 and 290 side and top views, respectively) and 482 narrow junctions (333 side views, 149 top views) (**Table S1**). The averages reached a resolution of ∼18 Å for the wide channel, and ∼26 Å for the narrow channel (**Fig. S4**). The wider channel had a pronounced pore of 2.7 nm throughout the entire length (**Fig. 3A, C**), while the narrow type had a pore of only 2 nm at the center, and appeared to be almost closed at the top (i.e. the protein densities at the respective cytosolic ends appeared to be in direct contact, with the pore possibly as narrow as 0.6 nm - though this is difficult to determine given the resolution; **Fig. 3B, D**). We did not observe protein density attributable to GFP, C-terminally fused to UNC-9. However we used animals that express UNC-9::GFP in an *unc-9* wild type background, i.e. GFP is present only on some copies of UNC-9, and may further be averaged out due to its flexible attachment. Both types of channels spanned both membranes and displayed an identical 5 nm membrane-to-membrane distance. Based on previous reports, GJs can adapt open or closed conformations, which is structurally related to the position of the N-termini of the single GJ subunits (17). This suggests that our wide and narrow averages might correspond to GJs in open and closed conformation, respectively. The channel top views (290 wide and 149 narrow channels) appeared to be arranged in a predominantly hexagonal pattern (**Fig. S3**). This observation was confirmed in STAs of top views (**Fig. 3C, D**).

**Figure 3:**
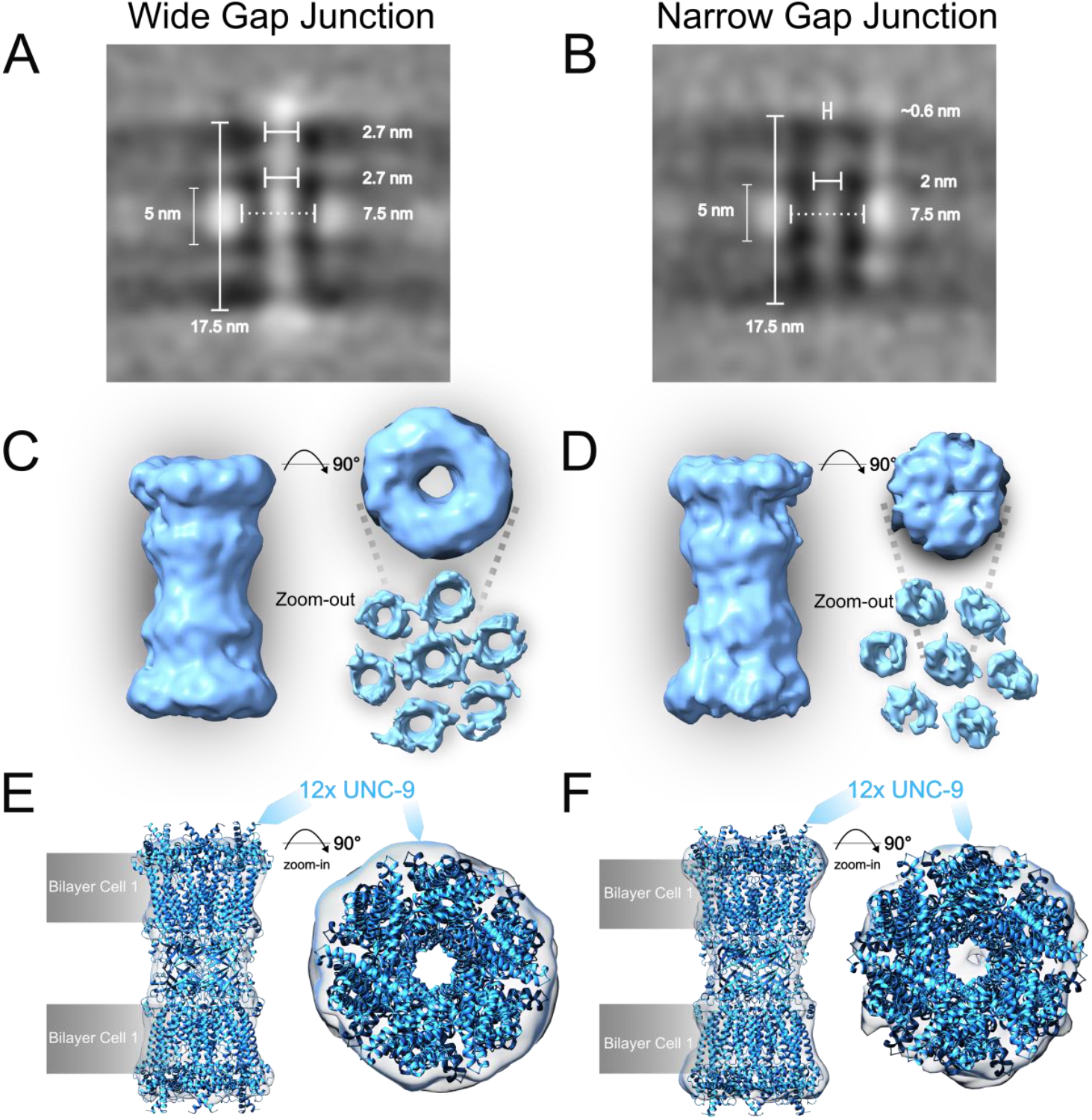
“Wide” (possibly open) and “narrow” (possibly closed) gap junction structures are observed following sub tomogram averaging and fitting of AF3 models. **(A, B)** 562 and 482 sub tomograms of wide, putatively open (A) and narrow, putatively closed GJ channels (B), respectively, were averaged to obtain a cryo-ET map. Dimensions are indicated. **(C, D)** Surface models, low-pass filtered to 30 Å, of the wide and narrow GJ channel, in side- and top-view, as indicated. Also shown are STA reconstructions with a widened mask, exposing the hexagonal array of six channels surrounding the central channel. **(E, F)** Structural model of the UNC-9 dodecameric innexon, derived from AlphaFold 3, are fitted into the *in situ* cryo-ET maps of the wide and narrow channels.

Given that structures of purified, detergent solubilized or nanodisc-reconstituted *C. elegans* GJs exhibited hexamers or octamers (16, 31-33), we next set out to determine the multimeric assembly of our *in situ* cryo-ET map. We used AlphaFold 3 to model different arrangements and combinations of the most abundant *C. elegans* innexins, i.e. UNC-7 and UNC-9 (**Fig. S5**). While the overall dimensions of the hexameric model (showing the highest overall confidence pIDDT value) appeared to match those of our average (**Fig. 3E, F; S5B**), the resolution of the STA was not sufficient to unambiguously determine whether hexameric or octameric assemblies were present *in situ*. The hexagonal arrangement in the GJ patches may render a hexameric channel assembly plausible. Yet, as the individual channels are quite distant to each other (∼3 nm), there may be no direct innexon-innexon contacts within the membrane. Thus, a hexagonal arrangement of octameric channels cannot be excluded, and examples for non-hexameric protein complexes forming hexagonal lattices have been observed previously (52, 53).

### Evidence for an unknown GJ type with a cytosolic cap structure

A subset of our tomograms exhibited additional structures on the cytosolic side in addition to the channel-like density extending across the two adjacent membranes (**Fig. 4A-F**). We found such structures in 20% of the GJs, and among those, there could be such a structure on only one side of the junction (76%), or on both sides (24 %; **Table S2**). When analyzing these junction types by STA (320 particles, resulting in 23 Å resolution; **Figs. 4G; S4**), we confirmed that the channel-like density was of the same dimensions as the GJ channel we observed earlier (**Fig. 3A-D**). Note that as we included all particles, the average density map exhibits a clear structure on one side, and a more diffuse density on the other side of the junction (**Fig. 4G**).

**Figure 4:**
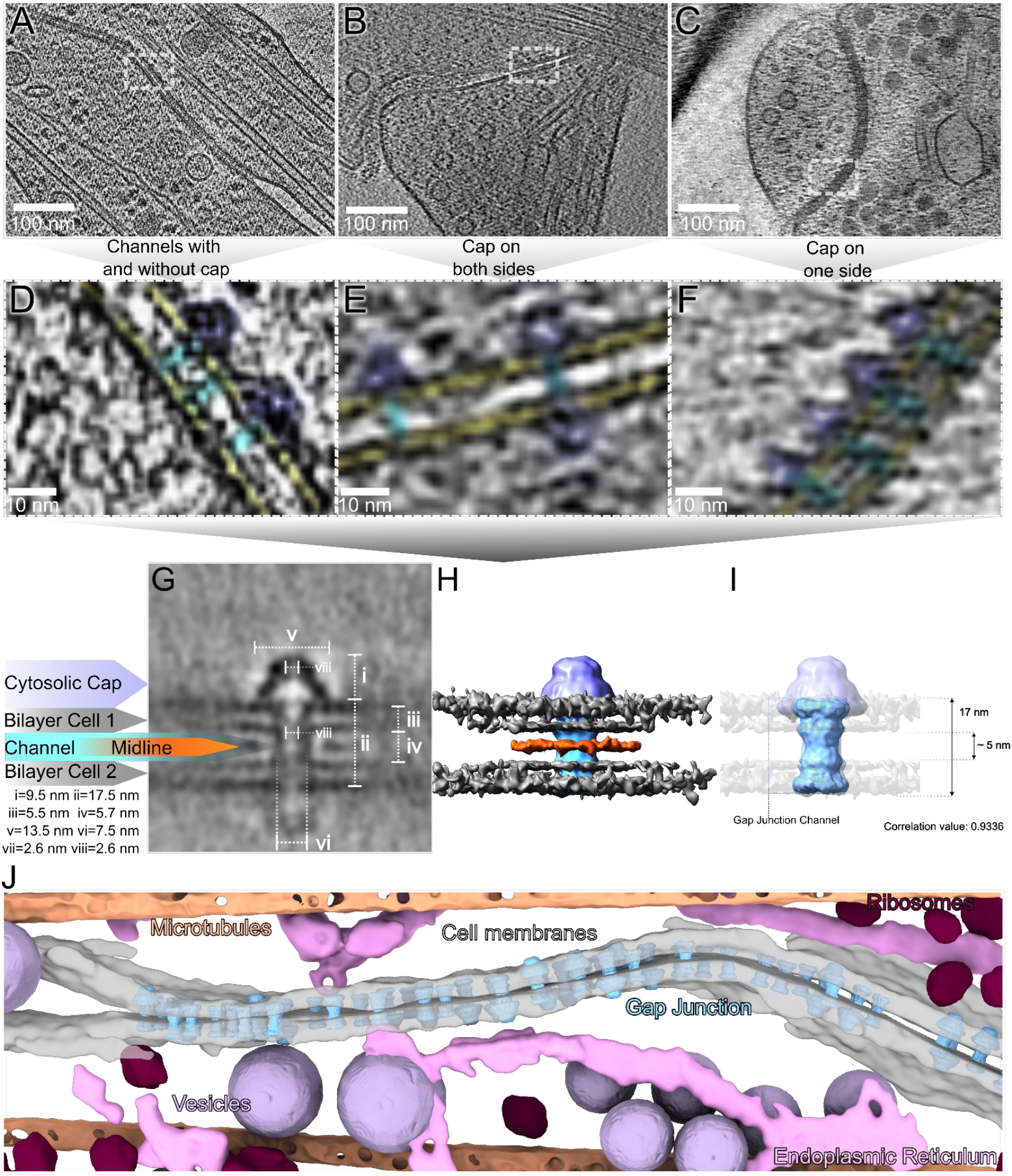
A subset of gap junctions exhibits a cytosolic cap structure *in situ*. **(A-C)** Example tomogram slices of gap junctions that expose cytosolic extensions. **(D-F)** Close-ups of the cell-cell contacts in (A-C), exhibiting a triangular cap structure on one or on both sides of the junction. Membranes are highlighted in yellow, GJs in cyan, and cap structures in dark blue. **(G)** STA reconstruction of 320 particles of the cap structure, showing its dimensions and hollow structure. Structural features are indicated. **(H)** 3D rendering of the surface map, low-pass filtered to 28 Å. The cap is shown in blue, the channel in cyan. **(I)** Fitting the surface map of the cap-less GJs (as shown in **Fig. 3C, D**), into the surface map of the ‘cap junction’; dimensions are indicated **(J)** Segmentation of cell membranes and organelles, including capped and uncapped GJs. See also **Movie S1**. Scale bars are indicated in each panel.

The additional structure seemed to reside on the cytosolic surface of the plasma membrane, centrally topping the intercellular channel. It resembled a cone-shaped, hollow cap of around 13.5 nm width at the base, and a thinner top, extending around 9.5 nm into the cytosol, with a ca. 2.6 nm pore; **Fig. 4G, H**). We will thus refer to it as cytosolic ‘cap’ in the remainder of the paper. To confirm that the capped channels are a type of GJ, we assessed the correlation of the cryo-ET map of the uncapped GJs with the channel of the ‘cap junction’ (**Fig. 4I**). Maps fitted well with correlation coefficients of 0.9336 and 0.8879 for wide/open GJ and narrow/closed GJ, respectively. The intercellular channels with a length of ∼17 nm and a ∼5 nm gap between cells were similar in both structures (**Figs. 3A, B; 4G-I**). Also, the channel widths and pore sizes of the channels were similar, with the dimensions of the capped channels being close to that of the wide GJ channels obtained earlier, with no cap structure attached (ca. 2.7 nm; **Figs. 3C; 4G-I**). Whether these GJ proteins are identical in each case is currently unknown.

In the STA, we noticed a dense midline between the cell membranes of the capped GJs, which we could not further identify (**Fig. 4G, H**, labeled in orange). The STA showed no direct connection of this midline to the GJ channels. This structure could be an extracellular extension of the cytosolic cap, i.e. if this is constituted from a transmembrane protein. However, due to the low particle number, evidence for this is currently lacking.

Some of our collected tomograms only contained either capped or uncapped channels, however, in tomograms containing both channel types, they appeared heterogeneously distributed rather than spatially clustered. Following segmentation of a tomogram containing a GJ patch, we fitted the surface models of GJs and capped GJs into the tomogram, revealing how these channels are arranged *in situ* (**Fig. 4J; Movie S1**).

### UNC-1 stomatin as the prime candidate for constituting the cytosolic cap

To identify the constituents of the cap complex, we focused on previous findings indicating that UNC-1 is essential for the function of UNC-9 containing gap junctions, and for their physical interaction, as shown by bimolecular fluorescence complementation assays (34, 35). The latter showed an interaction of the two proteins in close proximity, that required the C-terminal half of UNC-1 and the UNC-9 C-terminus. In an attempt to confirm the involvement of UNC-1 in capped junctions, we used a strain expressing UNC-1 C-terminally fused with GFP, i.e. supposing that an additional density might be observable. However, we did not find capped channels in cell cultures originating from this strain. Animals expressing UNC-1::GFP had locomotion phenotypes, just as mutants of the UNC-9 innexin (**Fig. S6**). Thus, possibly, UNC-1::GFP cannot properly assemble with UNC-9 GJs. However, we also tested a strain in which UNC-1 was N-terminally tagged with GFP. These animals showed no behavioral defects, and thus we used the CLEM approach to assess whether fluorescence would guide us to positions in the cell culture that show capped gap junctions. Indeed, such structures were observed (**Fig. S7**), emphasizing that UNC-1 is the likely constituent of the cytosolic cap. As an additional demonstration that UNC-1 can form the observed capped gap junctions along with UNC-9, we expressed the two proteins in a heterologous system, i.e. HEK cells. Also here, we could observe the capped gap junction structure in contact regions between two cells, spanning the two cell membranes (**Fig. S8**).

UNC-1 belongs to the SPFH protein family (**Fig. S9**), which includes also mammalian flotillin-1/-2 and bacterial HflK/C proteins. Recently, biological structures of these proteins were described, which resemble cap-like multimeric assemblies. Hence, we used AF3 structure predictions of UNC-1 (48), to investigate whether its multimeric assemblies (**Fig. S10**) match the dome-like entities observed in our *in situ* structures (**Fig. 4G-I**). Using eight or less monomers, AF3 could not model a circular assembly. However, when using nine or more subunits, AF3 predicted UNC-1 as cap-like structures (**Fig. S10**). The hexadecamer appeared to best match the dimensions of the cytosolic cap we observed *in situ* (**Figs. 5A; S11**). We used ChimeraX to fit the 16-mer of UNC-1 along with a 12-mer of UNC-9 (assembled from two hexameric hemichannels) into the cryo-ET map we generated from the ‘cap’ STA (**Fig. 5B-D**). The SPFH domain of UNC-1 would thus form the base, with the N-terminal α-helix possibly traversing the plasma membrane, a feature AF3 cannot predict, while the α-helices of the C-terminal regions wind around each other and form the top part of the cap (**Figs. 5A, D; S12**).

**Figure 5:**
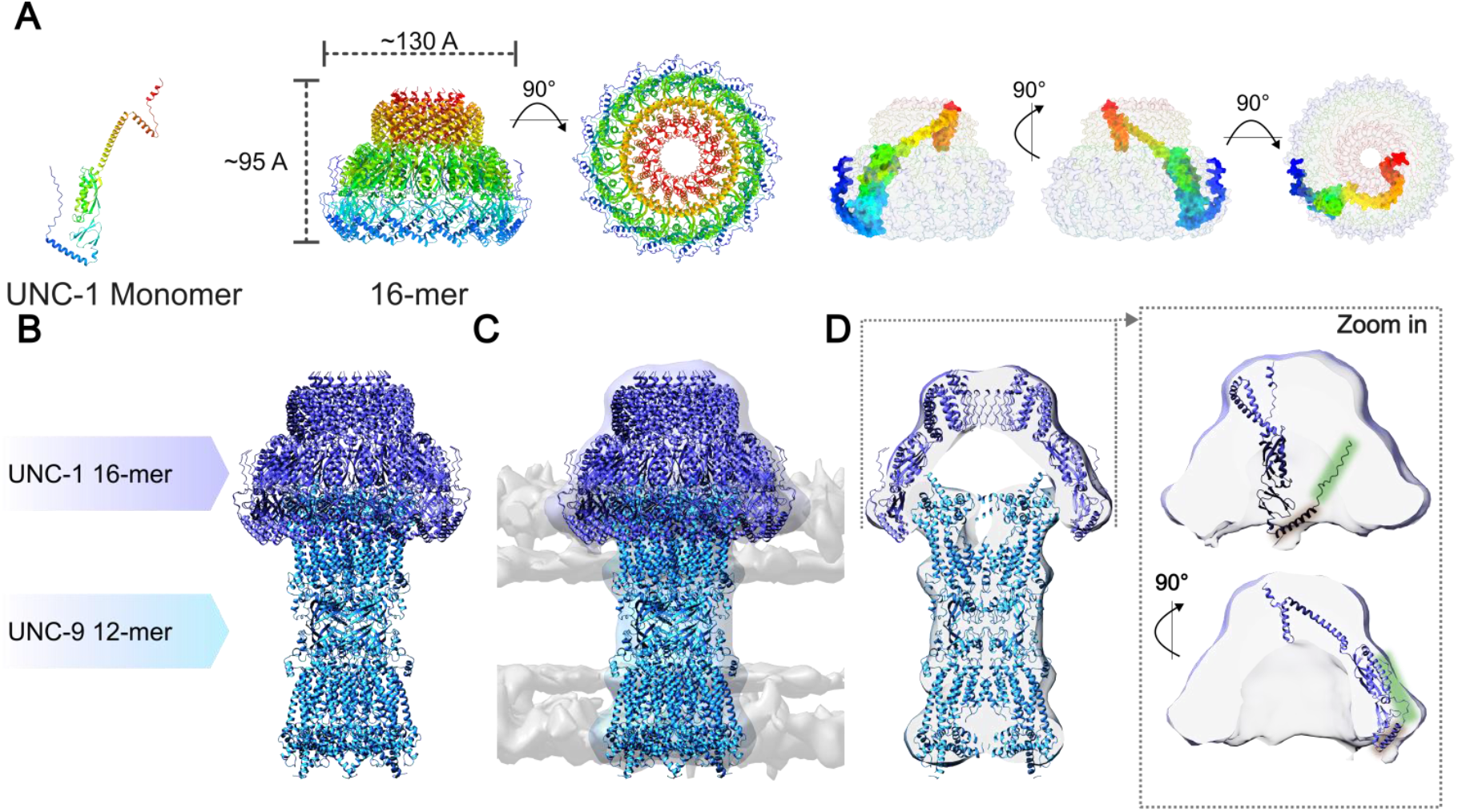
AF3 structure predictions suggest the cytosolic cap might be constituted by an UNC-1 stomatin multimer. **(A)** AF3 structural models of the UNC-1 stomatin. Monomeric structure as well as a hexadecameric ring model are rainbow color-coded from N-(blue) to C-termini (red). Also shown are the dimensions of the model and how single monomers fit into it. **(B)** Manual assembly and **(C)** fitting of the AF3 structural models of the UNC-1 hexadecamer (dark blue) and the UNC-9 innexin dodecamer (cyan) into the surface map of the ‘cap junction’. **(D)** Cross section through the model in (C) showing the close fit of the structural models to the *in situ* surface map of the capped gap junction. Close-up shows the fitting of a single UNC-1 model into the surface map. Note that the N-terminal α-helix of UNC-1 (brown shade in D, insert) is likely traversing the membrane, a feature that AF3 cannot predict. The unstructured N-terminus (green shade in D, insert) may thus reside outside of the cell.

Bacterial HflK/C and human flotillin protein complexes were shown to interact with the plasma membrane or other cellular membranes (flotillins, via lipid anchors attached to cysteins in the N-terminal SPFH1 domain (44)) or were inserted into the bacterial outer membrane (HflK/C, via an N-terminal extension with a membrane-traversing TM helix (45, 46)). Such assemblies might form a separated space, e.g. for chemical reactions, and/or to possibly confine or modulate the activity of transmembrane channels (44). UNC-1 resembles both flotillins and HflK/C in that it has an N-terminal hydrophobic (TM) helix as well as an SPFH1 domain with cysteins (**Fig. S12A**), and may thus interact with the plasma membrane both through (putative) membrane anchors, as well as a TM helix inserted into the membrane (**Fig. S12B-E**).

Also based on their surface potentials, the two structural models (UNC-1 hexadecamer and UNC-9 dodecamer), despite a symmetry break, appear to fit well, since negatively charged regions of the UNC-9 cytosolic surface would possibly meet positively charged regions on the concave side of the UNC-1 cap (**Fig. S13A**). Given the experimental evidence of UNC-9/UNC-1 interactions (34, 35) and the resemblance of the structural model with the density we obtained from cryo-ET, we suggest that the cap is indeed formed by the UNC-1 protein.

## Discussion

Gap junctions play a crucial role in neuronal networks by facilitating rapid signal transmission. Here, we determined the *in situ* structure of a gap junction from *C. elegans* primary cells and identified a complex of a GJ channel with an additional cytosolic protein structure. Given the morphology and dimensions of the protein density, the characteristic distance of the two adjacent cell membranes, and the hexagonal array of protein complexes, there is little doubt that the observed structures are gap junction channels. Fitting the molecular structures of the UNC-9 or UNC-7 GJs as obtained by AF3 modeling into the cryo-ET map was plausible and led to cross-correlation coefficients of >0.8. GJs in likely open or closed conformations were observed in equal proportions. The additional cytosolic, dome-like ‘cap’ structure is very likely to be formed by the UNC-1 stomatin, and the AF3-predicted hexadecameric UNC-1 cap structure fits the observed STA cryo-ET map very well. Our *in situ* structure currently lacks the resolution to unambiguously fit UNC-9 or UNC-7 innexins into the observed cryo-ET density map, and the identity of these innexins can only be assumed from the fact that GFP-labeling led us to find GJs in tomograms, while this does not exclude the presence of other innexins in these GJs. Whether the two classes of GJs we found are indeed open and closed cannot be firmly concluded at the obtained resolution. For example, the pores may have different sizes due to different innexin constituents However, in the ‘narrow’ GJ, the cytosolic ends of the densities appear to be in contact across the pore, and thus may indeed be closed. Also, the identity of UNC-1 as the constituent of the cap is inferred from indirect evidence, like the demonstrated functional and physical interaction, as well as the strong resemblance of the cap density with the AF3 model of the UNC-1 hexadecamer. However, the fact that we find capped gap junctions at sites in cell cultures labeled with GFP::UNC-1, as well as the finding of capped gap junctions in HEK cells expressing UNC-1 and UNC-9 lends further support to this hypothesis.

The functional meaning of this arrangement is currently unclear. UNC-9 was shown to localize to membranes in *unc-1(lf)* mutants (35), thus UNC-1 may not be required for trafficking and assembly of GJs. Yet, other functional roles for the UNC-1/GJ assembly are conceivable: 1) The UNC-1 cap could assist in GJ gating, by providing a binding site for the N-termini of the innexins. Some innexins like UNC-7b, but also UNC-9, have positive charges and, based on AF3 modeling, fold into the pore of the GJ, thus closing it. This mechanism was also proposed to underlie the closing of GJs in response to trans-junctional voltage differences (17). Once the N-termini are removed from the pore and thus unplug the GJ, they may need to be ‘parked’ elsewhere, such that they do not randomly close the pore in the absence of a voltage difference. Possibly, UNC-1 caps provide such a binding site (**Fig. S13B**). Looking at the surface potentials of the UNC-1 cap model, its inside exhibits rings of positive and negative potential (**Fig. S13A**). These might serve to provide binding interactions with the cytosolic surface of the UNC-9 complex, which exposes negative charges, but also with the positively charged innexin N-termini. 2) UNC-1 cages could serve as a selectivity filter for small molecules like 2^nd^ messengers and ATP (54). Such molecules, particularly cGMP, can be transferred from one cell to another through GJs (7). 3) The UNC-1 cap may help in gating the GJ, keeping the channel open (given that the cap channels resembled the open GJs), or in gating for particularly large cargoes. 4) The UNC-1 cap could help in the assembly of GJ channels, and UNC-1 rings may also assist in the formation of GJ patches in the membrane, or when individual channels need to be replaced. UNC-1 could then act as a placeholder for GJs in which one innexon is replaced by another. This might enable plasticity in electrical synapses. 5) The asymmetry imposed by binding of a cap (most of the observed capped GJs had a cap only on one side of the channel) could suggest a role in conveying rectification properties. Heterotypic GJs made from UNC-9 and the UNC-7b splice variant exhibit rectification properties in *Xenopus* oocytes (11). UNC-1 was not included in these experiments, thus rectification may not require UNC-1 for this combination of innexin subunits. However, UNC-1 could convey rectification properties to homotypic GJs by binding from one side only. Such hypotheses may be addressed now with specific experiments, e.g. once a more precise interaction surface of UNC-1 caps and innexin GJs can be established, for example by molecular dynamics simulations.

The details of the structural arrangement of UNC-1 at the membrane are currently unclear. UNC-1 contains an N-terminal α-helix like the bacterial HflK/C proteins, which in the biological structure traverse the membrane, and it has cysteins in the SPFH1 domain, like flotillins, which penetrate the outer leaflet of the membrane and are palmytoylated, acting as membrane anchors. UNC-1 may thus also insert its N-terminal TM helix into (**Fig. S12E, F**), while the attachment through the SPFH1 domain may assist in assembling the UNC-1 cap at, the plasma membrane. For the UNC-1 homologue MEC-2, analogous cysteins were shown to be palmytoylated, and the proline inducing the bend in the N-terminal α-helix (P48 in UNC-1; **Fig. S12A, B**), when mutated to serine, affected MEC-2 membrane topology (55, 56). If the N-terminal helix of UNC-1 is inserted into the PM, this would place the N-terminal ca. 30 amino acids outside the cell, possibly constituting the dense midline we observed in association with the capped junctions (**Figs. 4G, H; S12F**).

Our work sheds new light on GJ *in situ* structures and their assembly with an associated protein complex. Since stomatins are found in all animals, it is likely that similar assemblies exist also in vertebrates. Candidate UNC-1 homologues are human STOM and podocin. Stomatins were associated with the function of other ion channels, e.g., mechanosensory DEG/ENaC channels require the stomatin MEC-2 (57, 58). Here, a role in the organization of the mechanosensory channel by MEC-2-mediated phase separation was uncovered. Stomatin functions have also been associated with ion conductance in red blood cells *via* acid-sensing ion channels (ASICs, related to ENaCs), glucose transporters, or TRPC channels (59). In light of this evolutionary conservation and diversity of roles, a function for stomatin caps also in the context of mammalian gap junctions appears possible.

## Materials and Methods

### Molecular Biology

To express the respective marker proteins in *C. elegans* transgenic strains, we used the p*unc-17* (cholinergic neurons), p*rgef-1* (pan-neuronal) and p*rab-3* (pan-neuronal) promoters. As a selection marker we used mCherry expressed under the p*myo-2* (pharyngeal muscles) promoter. The plasmid pNR13 [prgef-1::UNC-9::GFP] was assembled as follows: The p*rgef-1* promoter sequence was amplified from pJH4531 [prgef-1::DAF-2] (Addgene plasmid #132366) using primers oNR016 (5’-GAAATAAGCTTGGGCTGCAAGCTTGCATGCCTGCAGCG-3’) and oNR017 (5’-GCATACTCATCCTGGATCTTTACTGCTGATCGTCGTCGTCG-3’). The amplicon was ligated into vector pNR07/p612 (a gift from Zhao-Wen Wang (12)) following restriction digestion with PstI and BamHI by Gibson assembly. Further plasmids used were p1676[punc-17(short)::tagRFP::ELKS-1] (a gift from Zhao-Wen Wang), pFR227[prab-3::rab-3::mCherry], pAB20[punc-17::ChR2(H134R)::mTFP] and pAG8[plev-1::lev-1-HA::6xHIS-3xHA] (60).

### *C. elegans* strains

*C. elegans* strains were cultivated according to standard methods (61). Animals were kept on nematode growth medium (NGM), and fed *Escherichia coli* (*E. coli*) strain OP-50-1, as well as *E. coli* strain Na22 for cell culture experiments (see *C. elegans* primary cell culture). Transgenic strains were generated by microinjection of DNA plasmids into the gonads (62) of either wildtype (Bristol N2) or juSi164 (Histone-miniSoG) animals. Integration into Histone-miniSOG strain juSi164 was performed as previously described (63).

Strains used or generated: **Bristol N2, CB719:** *unc-1(e719)X*, **CZ20310:** *unc-119(ed3) III; juSi164[mex-5p::HIS-72::miniSOG; Cbr-unc-119(+)]*, **MT1088:** *unc-1(n494)X*, **OH16682:** *unc-1(ot1076[UNC-1::GFP])*, **ZX914**: *unc-9(e101)*, **ZX3070**: *zxIs148[punc-17(short)::tagRFP-ELKS-1; pmyo-2::m-Cherry]*, **ZX3773/OH15255:** *unc-7(ot895[unc-7::tagRFP])X*, **ZX3776**: *unc-1(ot1076[UNC-1::GFP])* **ZX3905**: *zxIs266[prgef-1::unc-9::GFP]*, **ZX3906**: *unc-7(ot895[unc-7::tagRFP]); unc-1(ot1076[UNC-1::GFP])*, **ZX3973**: *zxIs288[prab-3::rab-3::mCherry; punc-17::ChR2(H134R)::mTFP; plev-1::lev-1-HA::6xHIS-3xHA]*.

### CRISPR/Cas9-mediated genome editing

C-terminal GFP::3XFlag-tagged knock-in allele of *unc-1* was generated using CRISPR/Cas9-mediated homologous recombination based on (64); crRNA sequence: 5’-aaaaaggaaaatatgaTTAT-3’.

### Behavioral assays

For the analysis of crawling and swimming behavior, 30 to 80 L4 animals were picked onto a fresh NGM plate with a small lawn of OP-50 bacteria in the center, the day before the measurement. For crawling assays, these plates were directly transferred to the multi worm tracker platform (65) equipped with a high resolution camera (Falcon 4M30, DALSA) and a custom-built infrared transmission light source (6WEPIR3-S1, WINGER, 850nm 3W). Videos of the animals were recorded for 10 minutes. Kink was extracted using Choreography (65). Bulk processing of multiple files for Choreography was handled with a custom-written python script (https://github.com/dvettkoe/MWT_Analysis). For swimming assays, animals were washed off the NMG-plates with OP-50 and after three washing steps transferred to fresh, 3.5 cm NGM plates without bacteria. Worms were kept swimming on these plates in 800 μL M9 buffer for 10 to 15 minutes before being placed on the multi worm tracker platform. 1 minute videos of swimming behavior were acquired, using a LabVIEW-based custom software (MS-Acqu) (66). Swimming cycles were analyzed with the “wrMtrck” plugin (67) for ImageJ. The automatically generated tracks were validated with a custom written Python script (https://github.com/dvettkoe/SwimmingTracksProcessing).

### Data and statistical analysis of behavioral assays

Data are shown as median with interquartile range, n indicates the number of animals, and N the number of biological replicates (experiments with different sets of animals). Significance between datasets after one-way ANOVA with Bonferroni’s multiple comparison test is given as a p-value. Data were analyzed and plotted in GraphPad Prism (GraphPad Software, version 8.02).

### *C. elegans* primary cell culture on EM grids

Primary cell culture was performed as described by (68) with a few improvements as well as adaptations to cell culture on EM grids. Nematodes were grown on Na22 *E. coli* plates until late adulthood, when they contain a maximum of eggs. Next, animals were washed off the plate with 7 mL ddH_2_O and transferred to a 15mL Falcon tube. 2 mL 2.8 % w/v NaClO (household bleach) as well as 1 mL of 5M NaOH were added. The suspension was vortexed until the adult worms were fully dissolved. The remaining eggs were washed with sterile ddH_2_O (10 mL) under a cell culture hood for three times, while pelleting the eggs in between each wash at around 900 rcf (g). After the third wash, 0.5 mL Chitinase (C6137, Sigma; 1 unit / mL, in egg buffer) was added and eggs were transferred to a 1.5mL centrifuge tube. During a 1-1.5 hour period, eggs were rotated at RT. The chitinase reaction was stopped by adding 0.8 mL of L15 full medium (Thermo Fisher). Eggs were pelleted at 900 g in a tabletop centrifuge for 3 minutes. After removing the supernatant, 500 μL of L15 full medium were added and cells were dissociated. To this end, the solution was taken up into a 2 mL syringe with a 19-gauge needle and pushed out back into the centrifuge tube. This was repeated 15 to 20 times until embryos were fully dissociated. The state of dissociation was monitored under a light microscope at 40x magnification by pipetting 1 μL of the suspension on a microscope slide after 15 or 20 repetitions, respectively. After dissociation, four fresh centrifuge tubes were prepared, 1 to 4, the original tube will henceforth be named tube 0. 1 mL of L-15 full medium was added to tube 0, and taken up with the syringe. The needle was replaced with a 5 μm syringe filter and the solution was pushed through the filter into tube 1. The amount of liquid was adjusted by additional L15-medium to 1mL. After emptying the syringe, tube 0 was again filled with 1 mL L-15 full medium and the filter was again replaced by the needle. The solution was taken up again, the needle was exchanged for the filter and the cell solution was pushed through the filter into tube 2. This process was repeated for tubes 3 and 4. Cells were spun down at 900g for 5 minutes in a tabletop centrifuge. The supernatant was discarded carefully, leaving about 50 μL in each tube. With a 200 μL pipette, cells were resuspended and pooled in tube 1, starting from tube 4 to reduce the amount of cells lost during the process.

Carbon and gold bead coated EM grids (Lacey, Au Finder, 200 mesh) were glow-discharged and coated with peanut lectin for 20 minutes, washed with ddH_2_O and dried for 20 minutes. Grids were then placed in glass-bottom dishes and 300 to 800 μL of cell suspension was applied to the grids. Cells were allowed to adhere and differentiate for 16 to 24 hours before imaging.

### Fluorescence imaging of primary cell culture samples on EM grids

Primary cells expressing fluorescently labelled marker proteins were examined under an Axio ObserverZ1 (Zeiss). Images were recorded with Teledyne Photometrics Kinetix22 camera controlled with an ImageJ plugin MicroManager1.4 (ImageJ version v1.48, (69, 70)). The distinctive carbon pattern and labelling of lacey gold finder grids allowed fluorescently marked junctions to be registered for subsequent examination with cryo-TEM.

### Vitrification of primary cell cultures

Primary cells grown on EM grids, were vitrified via plunge freezing using the Vitrobot Mark IV (Thermo Fisher) and a back-sided blotting approach by which two blotting papers were mounted on the blot pad facing the back side of the grid, and a Teflon sheet was mounted on the blot pad facing the sample side of the grid. The chamber was then equilibrated at 60% humidity and 20 °C for 60 min before starting the experiment. Then, EM grids were retrieved from the culture dish and additional 3 μl of cell culture medium were added to the grids prior to blotting. Finally, blotting and plunge freezing into liquid ethane was performed with the following parameters: Blot force -1, Blot time 2.5 s, single blot event, Drain time 0.5 s.

### Cryo-Light Microscopy

EM grids were imaged by confocal laser scanning microscopy on a LSM700 microscope (Zeiss) using a CMS196 cryo-stage (Linkam) at -195°C. Images were acquired using an excitation wavelengths of 488 nm and 555 nm for GFP and mCherry or RFP, and 639 nm for reflected light, respectively. Images were acquired with 5x/NA 0.16, 20x/NA 0.4 or 100x/NA 0.75 objectives. Montage maps were generated manually using the 20x magnified images that were stitched Fiji version 1.51w using the Pair-wise stitching plug-in (71).

### FIB-milling

The plunge-frozen EM grids were clipped into cryo-FIB autogrids (Thermo Fisher) and loaded into a pre-tilted EM grid holder with shutter and additional cold trap (Leica) under liquid nitrogen using an EM VCT500 loading station (Leica). Then the sample holder was taken up by the EM VCT500 manual transfer shuttle and transferred into the Leica EM ACE600 for platinum sputtering (4 nm). Afterwards the sample holder was transferred - via a VCT dock (Leica) – into the Helios 600i Nanolab FIB/SEM dual beam instrument equipped with a band-cooled cryo stage (Leica) equilibrated at -157 °C. The EM grids were imaged using SEM (3 kV, 0.69 nA) and FIB (gallium ion source, 30 kV, 33 pA). An overview SEM image was acquired at 90° incident angle, followed by 2D correlation of SEM and cryo-LM images with ImageJ using the Bigwarp plug-in (72). As fiducials for the 2D correlation, features in the carbon film, such as lacey carbon strands, were used. *C. elegans* primary cells were chosen for milling at a net incident angle of 12°. First, trenching was done at 240 to 400 pA (FIB), then a few μm thick organometallic Pt layer was deposited (GIS valve open for between 10 to 30 s). A first thinning step was performed with 400 pA down to a pre-lamella thickness of ∼1.5 μm. A second thinning step was done with 83 to 240 pA down to a pre-lamella thickness of ∼0.4 μm. The final lamella thickness of around 0.2 μm was reached by a last thinning/polishing step at 33 pA.

### Cryo-TEM imaging and Cryo-Electron Tomography

Movie-stacks of thin, plunge-frozen primary cells were imaged at a Titan Krios cryo-TEM (FEI) equipped with a field emission-gun operating at 300 kV, a GIF Quantum post-column energy filter (Gatan) operated at zero-loss and a K2 Summit or a K3 direct electron detector (Gatan). Low magnification montages were acquired at a nominal magnification of x11,500 at 0° stage tilt or at the lamella pre-tilt angle induced by FIB milling (corresponding to 12.56 Å/px on the K2 and 8.22 Å/px on the K3) and a defocus of -50 μm with a total electron dose of 0.5 e^-^/Å^2^ using SerialEM (v3.8 and v4.1) (73). Dose-fractionated tilt series were acquired at a nominal magnification of ×64,000 (corresponding to 1.1 Å/px on the K2 and 0.72 Å/px on the K3) using the SerialEM low dose acquisition in super resolution mode. A dose-symmetric tilt-scheme with an angular increment of 3° (−60° to +60°, with respect to the lamella pretilt if applicable) with a cumulative dose per tilt series ranging between 145 e^-^/Å^2^ and 150 e^-^/Å^2^ and a defocus of - 3 μm.

### Image processing / Data analyses

#### Tomogram reconstruction

Frame motion was corrected using ImageStackAlignator (https://github.com/kunzmi/ImageStackAlignator, (74)) and tilt series were aligned using patch tracking in IMOD (version 4.9.12) (75) and reconstructed through weighted back projection in ARTIATOMI (https://github.com/uermel/Artiatomi) (76) using the IMOD alignment. For visualization purposes, tomographic reconstructions were low pass filtered in Fourier space with a cut-off at 1 nm, and the contrast was enhanced by applying a Wiener-like deconvolution filter (https://github.com/dtegunov/tom_deconv). For sub-tomogram averaging sub-tomograms were reconstructed by weighted back projection with exact filter and contrast transfer function (ctf) correction. The ctf was estimated using CTFFIND4 v4.1.13. (77). Particle picking and pre-orientation relative to the membrane was done in ArtiaX (78). All particles were randomized in Phi (i.e. angle around the z-axis of the sub-tomogram). Subtomogram averaging was done with ARTIATOMI wrapper for MATLAB 2024a (The MathWorks). For the gap junctions, particles were binned 4 times. A total of 562 and 482 particles were used for the final average of wide and narrow channels, respectively. For the cap junction, particles were also binned 4 times. A total of 320 particles were used for the final average. The pixel size of the data set with the larger pixel was kept constant (K2) and the relative pixel size of the second data set was adjusted (K3) to merge datasets of different pixel sizes. Those particles were scaled to a pixel size corresponding to 4.4 Å/px using MATLAB 2024a. Resolutions were estimated using the FSC 0.143 criterion. FSC plots were calculated in RELION-5 (http://www.github.com/3dem/relion (79)), using randomized half sets generated in ARTIATOMI. A customized mask was applied to both half-maps. The resolution of the averages was estimated at 18 Å and 26 Å for wide and narrow channels, respectively and 23 Å for capped junctions.

#### Fitting of published EM structures or AlphaFold predictions into STA-results

The structures of AlphaFold3-predicted models (48) of UNC-7, UNC-9 and UNC-1 were rigid-body fitted into our averages using ChimeraX v1.8 (71, 72). The fit quality was visually assessed.

#### 3D Segmentation

Automated segmentation was performed with Dragonfly (Version 2024.1 for Windows, non-commercial license. Comet Technologies Canada Inc., Montreal, Canada; available at https://www.theobjects.com/dragonfly) and membrane segmentation was performed with MemBrain-seg (teamtomo, https://github.com/teamtomo/membrain-seg) on deconvolution-filtered tomograms. The smoothness of respective segmentations was improved using mean curvature motion ((80); (https://github.com/FrangakisLab/mcm-cryoet). The resulting segmentation maps were animated in the ArtiaX tool implemented in ChimeraX v1.6 (https://github.com/FrangakisLab/ArtiaX).

## Acknowledgments

We thank Katharina Kuhlmeier, Franziska Baumbach, and Hans-Werner Müller for expert technical assistance and Oliver Hobert, Zhao-Wen Wang and Mei Zhen for providing reagents. Furthermore, we thank Margot P. Scheffer, Mbuso S. Mantanya, Lisa Rehm und Deborah Moser for assisting with computational experiments run in CryoSPARC, RELION-5 and Dragonfly. We are indebted to members of the Gottschalk and Frangakis labs for advice and critically reading the manuscript. Some strains were provided by the Caenorhabditis Genetics Center (CGC), which is funded by by NIH Office of Research Infrastructure Programs (P40 OD010440).

## Author contributions

Conceptualization: NR, KW, ASF, AG

Methodology: NR, KW, ANB, SM, ASF

Reagents: AB

Investigation: NR, KW, ASF

Visualization: NR, KW, ANB, ASF, AG

Funding Acquisition: ASF, AG

Project administration: ASF, AG

Supervision: ASF, AG

Writing – original draft: NR, AG

Writing – review & editing: NR, ANB, SM, ASF, AG

## Data and materials availability

The datasets generated during and/or analyzed during the current study are available from the corresponding authors on reasonable request. This also applies to materials described in the study. The cryo-ET structure solved in this study is available in the Electron Microscopy Data Bank (EMDB) under the accession EMD-XXXX.

## Supplementary Materials

Figures S1-S13

Tables S1, S2

Movie S1

