## Supplementary Figures and Tables for "*In situ* structure of a gap junction – stomatin complex"

Nils Rosenkranz *et al.*

**This PDF file includes:**

Figs. S1 to S11
Tables S1 to S2
Movie S1

**Other Supplementary Materials for this manuscript include the following:**
Movie S1

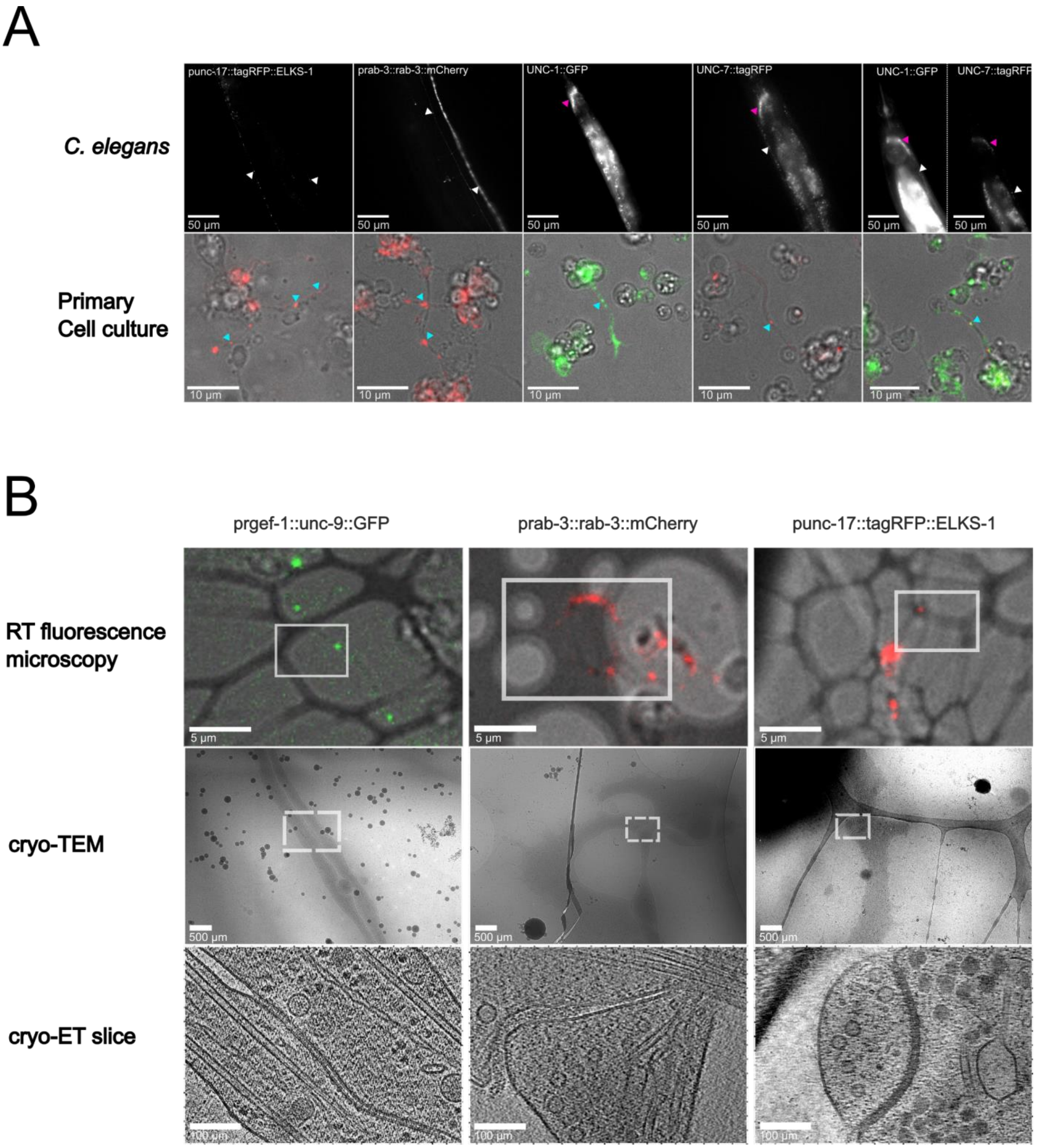


**Fig. S1. *C. elegans* strains used for the study of *in situ* cell-cell junctions. Cell cultures and example fluorescence imaging, electron microscopy and tomography. (A)** Fluorescent micrographs showing *C. elegans* (top row), or primary embryonal cell cultures derived from these animals (bottom row), expressing marker proteins, as indicated. Arrowheads point to synapses and gap junctions in nerve cord (white), nerve ring (magenta) and primary neuronal contact sites (cyan). **(B)** Example images of the indicated stages of the process of finding cellular junctions, by fluorescence microscopy, cryo-transmission electron microscopy (cryo-TEM), and cryo-electron tomography (cryo-ET), for the indicated transgenic strains, including gap junctions and possibly chemical synapses, as indicated. From top to bottom, boxes indicate the region shown in the next image as a close up. Scale bars are indicated in each panel.

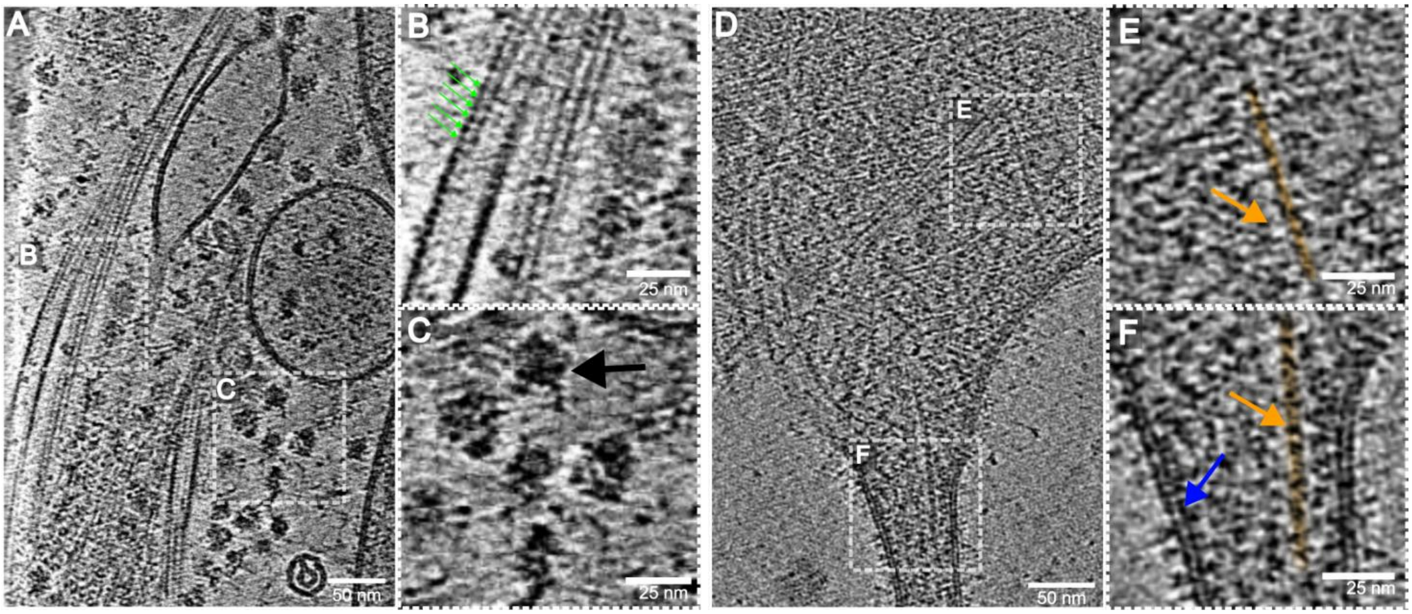


**Fig. S2. Cryo-ET characterization of *C. elegans* cytoskeleton and ribosomes in cell culture. (A)** Overview tomogram showing microtubule (enlarged in **(B)**; green arrows point at tubulin repeats) and ribosomes (enlarged in **(C)**; black arrow). **(D)** Overview tomogram showing actin filaments and cell membranes (enlarged in **(E)** and **(F)**; actin filament highlighted in orange and indicated by orange arrow; cell membranes indicated by blue arrow). Scale bars are indicated.

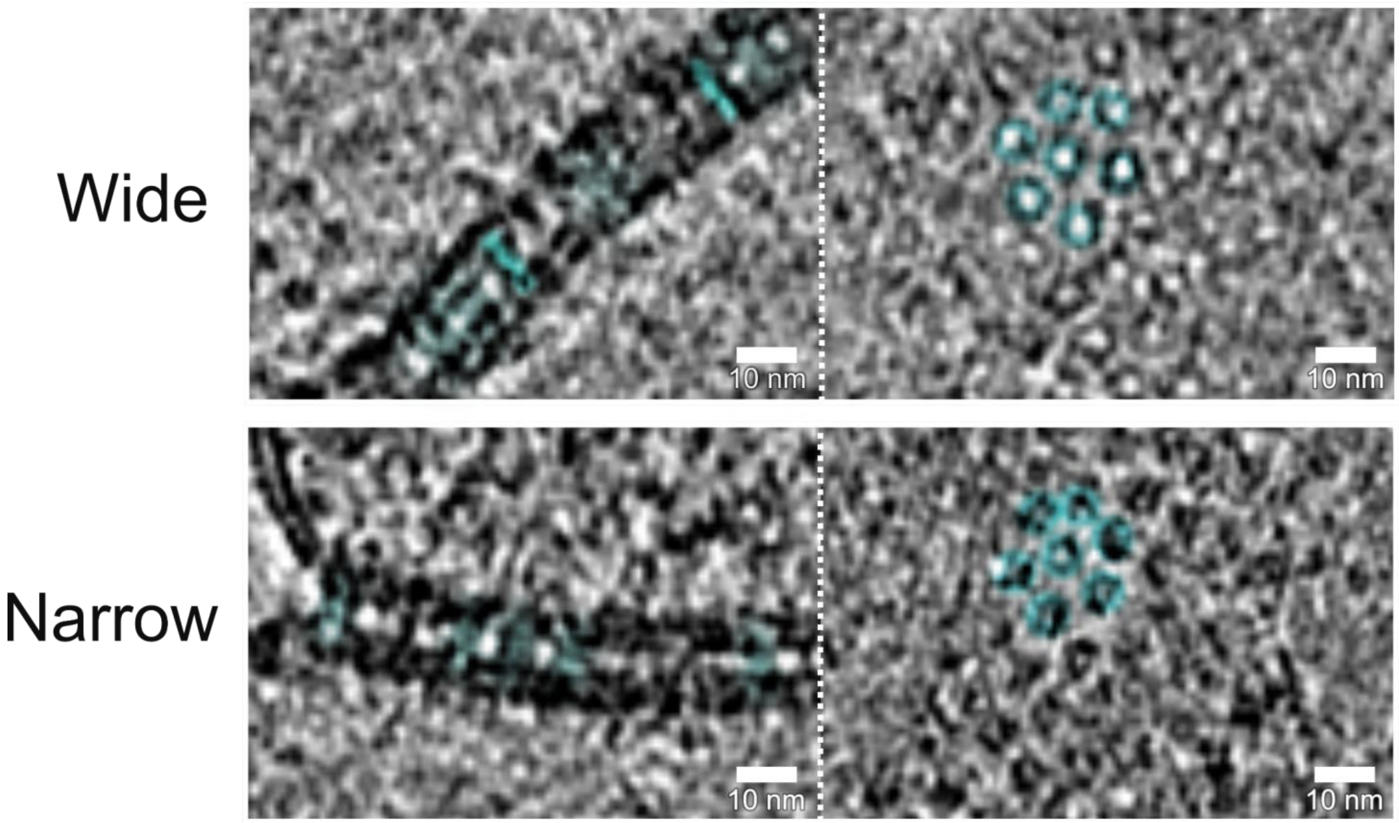


**Fig. S3. Cryo-ET images of open and closed GJ channels in side view, and in top view, in hexagonal arrangement. Top row:** Open GJ structures. **Bottom row:** Closed GJ structures. Scale bars are indicated.

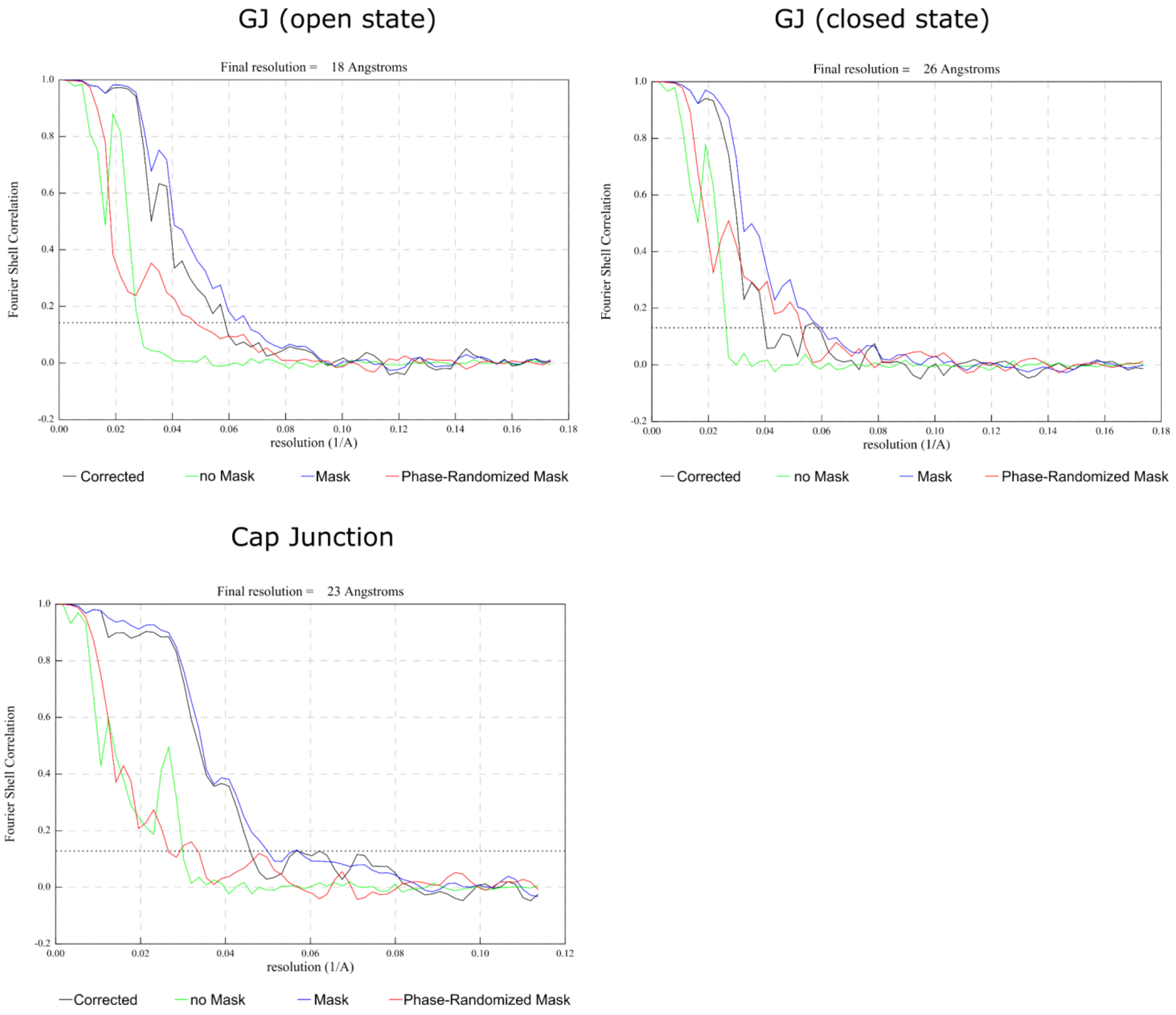


**Fig. S4. Fourier Shell correlations of open, closed and capped gap junctions.** Shown are Fourier Shell Correlation (FSC) curves of two half-datasets of the different subtomogram averages. The estimated resolution of the averages was 18 Å, 26 Å and 23 Å, for open, closed, and capped junctions, respectively. The resolution was calculated according to the gold standard criterion of FSC 0.143.

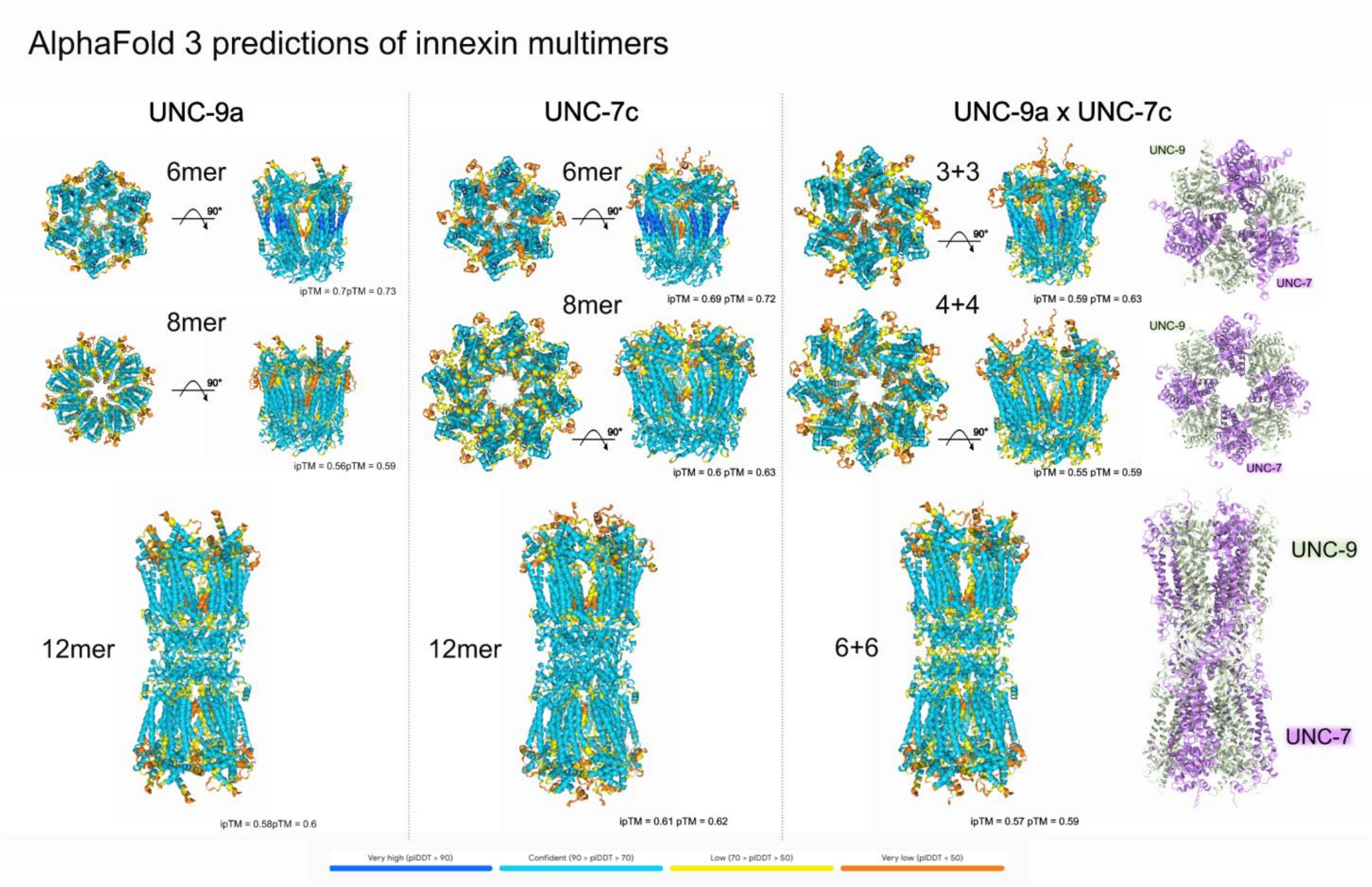


**Fig. S5. AlphaFold 3 structural models of abundant *C. elegans* innexins.** Homo-hexamers and octamers, as well as -dodecamers were modelled of the UNC-9a (left) and UNC-7c (middle) innexins. Also, heteromeric UNC-9a/UNC-7c GJ assemblies were modelled (right, as indicated). Confidence scores (ipTM - integrated predicted template modeling), and colour-coded plDDT (predicted local distance difference test, a per-residue confidence metric) values of the structural models are indicated.

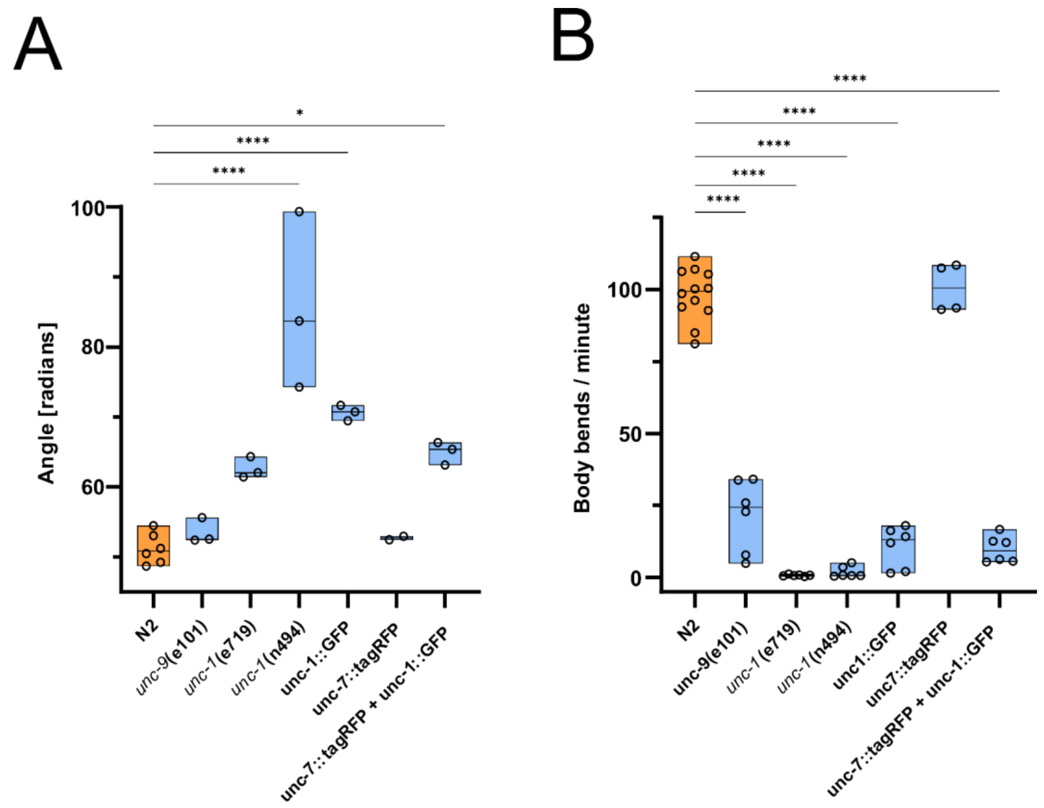


**Fig. S6. Locomotion behavior of different *C. elegans* strains analyzed in this work. (A)** Median body bending angles, as well as minimum and maximum values, of worms of the indicated genotypes were measured on NGM plates with OP-50 lawn. N = 2 to 6 experiments with n = 30 to 80 animals were analyzed. **(B)** Median swimming cycles of animals transferred to and measured in M9 buffer, scored as full body bends, are shown per minute. Shown are medians as well as minimum and maximum values. N = 4 to 12 experiments with n = 15 to 70 animals were analyzed.

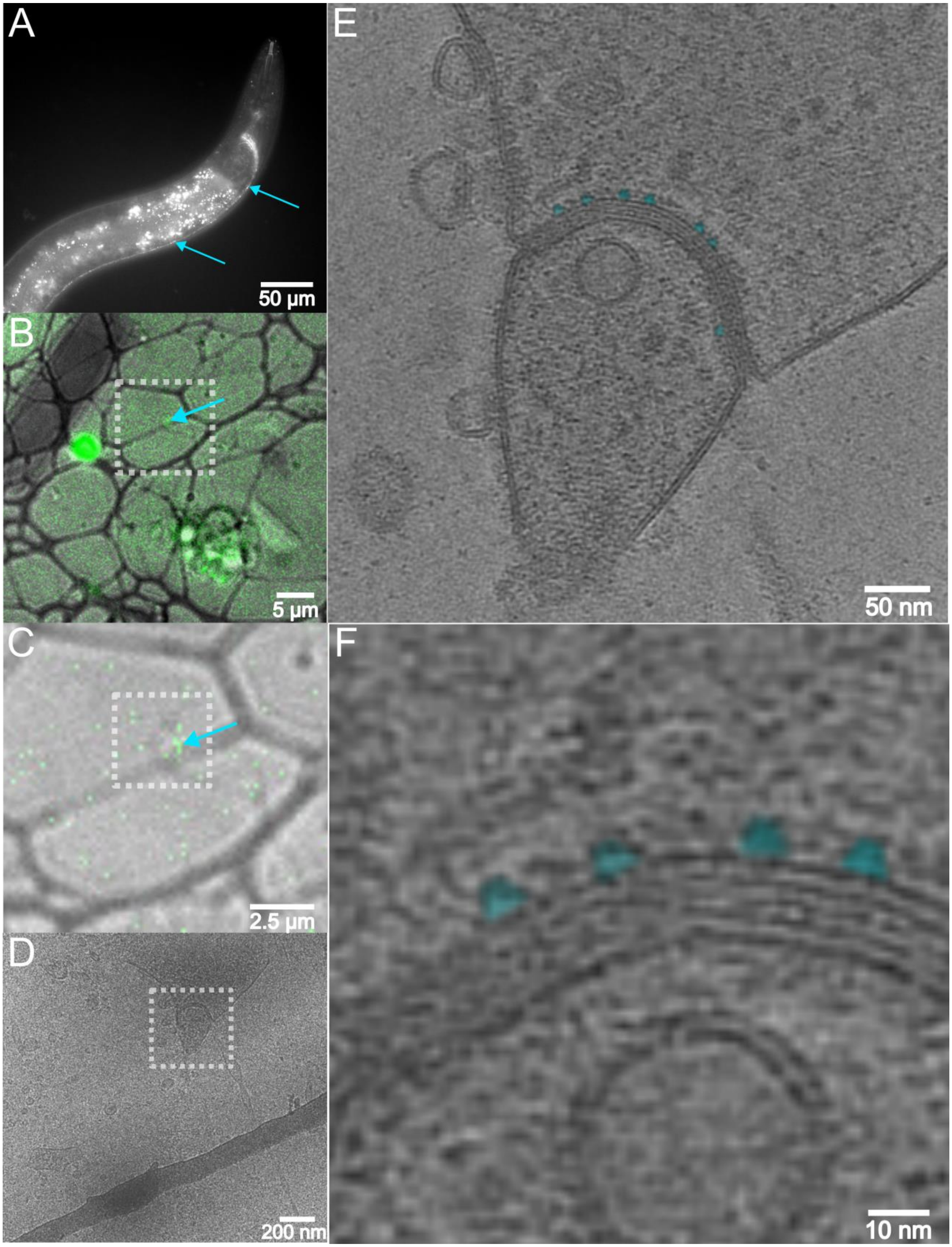


**Fig. S7. Capped gap junctions can be observed in cell cultures from a *C. elegans* strain expressing GFP::UNC-1. (A)** Animal expressing GFP::UNC-1 in the nervous system, in the nerve ring and along the ventral nerve cord (arrows). **(B-E)** CLEM approach leading to identification of a gap junction in cultured cells from the GFP::UNC-1 strain. **(F)** Zoom-in of a tomogram exhibiting the previously observed cap structures (shaded in cyan).

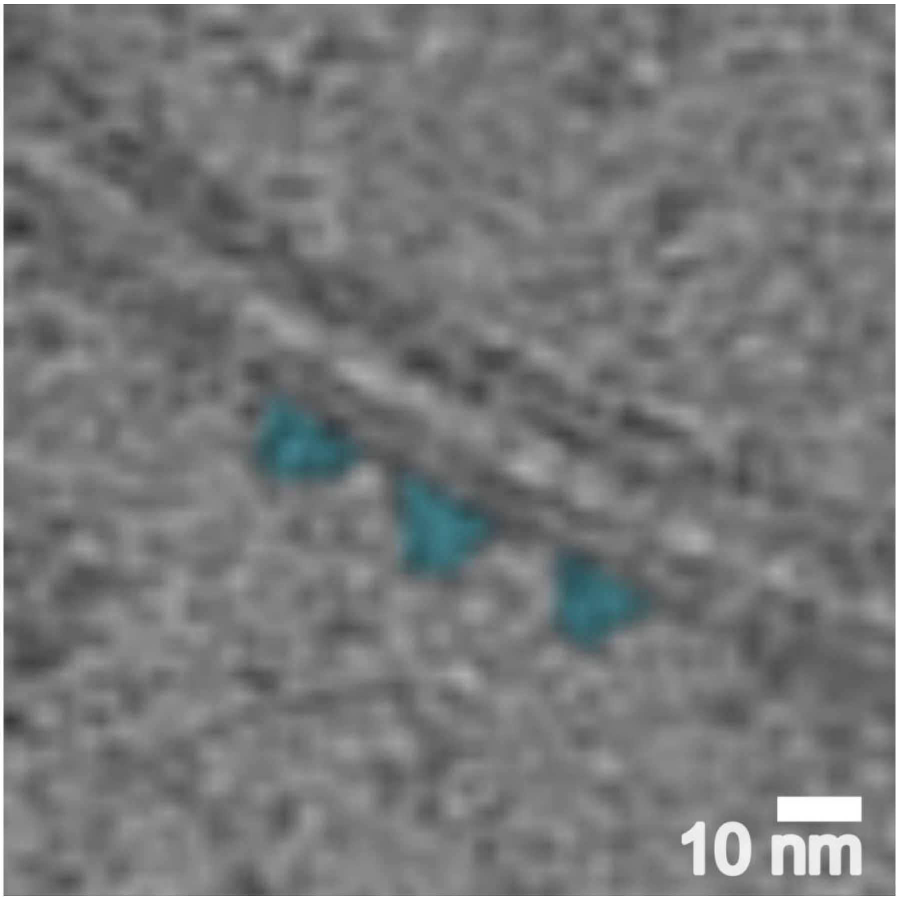


**Fig. S8. Capped gap junctions can be observed in heterologous cell cultures (HEK cells) expressing UNC-1 and UNC-9 cDNAs.** Close-up of a tomogram showing capped (cyan shade) membrane-spanning channel structures at a cell-cell contact of a distance typical for gap junctions.

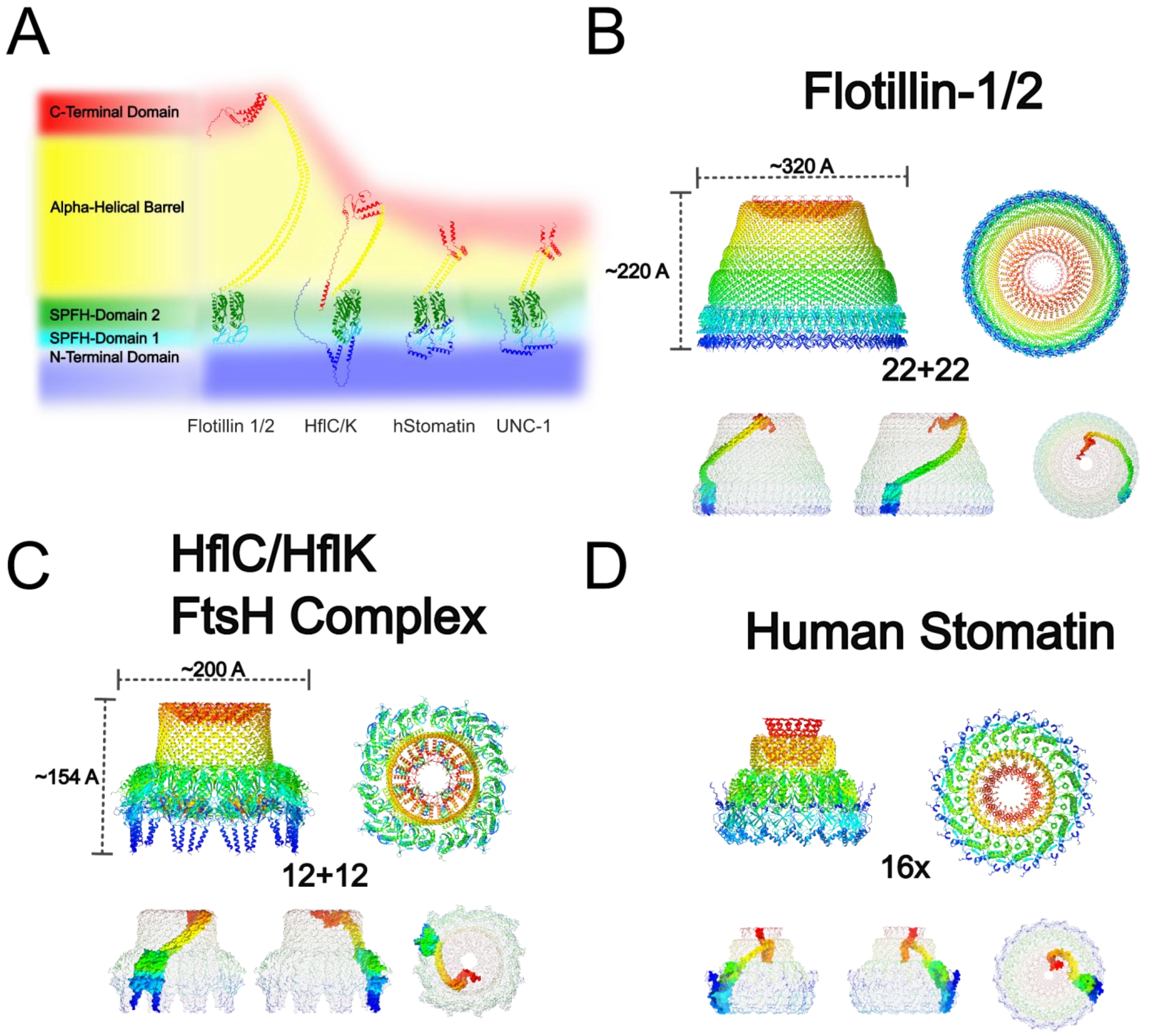


**Fig. S9. Proteins of the stomatin, prohibitin, flotillin and HflK/C (SPFH) domain family, AF3 structural models. (A)** Domain structure of the SPFH family is indicated and color coded, superimposed on AF3 models of the indicated single proteins. **(B-D)** Multimeric AF3 models of 22 + 22 subunits of human flotillins 1 and 2 (B); 12 + 12 bacterial HflKC proteins (C) and 16 copies of human stomatin (D). Colour code, rainbow from N- to C-termini, blue to red, resembles the domain annotations as in (A). Cryo-EM maps and atomic models downloaded from Electron Microscopy Data Bank (EMDB) and Protein Data Bank (PDB) with the following accession codes: Flotillin: EMD-44792 (EMDB) and 9BQ2 (PDB); HflC/K: EMDB-32002 (EMDB) and 7VHP (PDB).

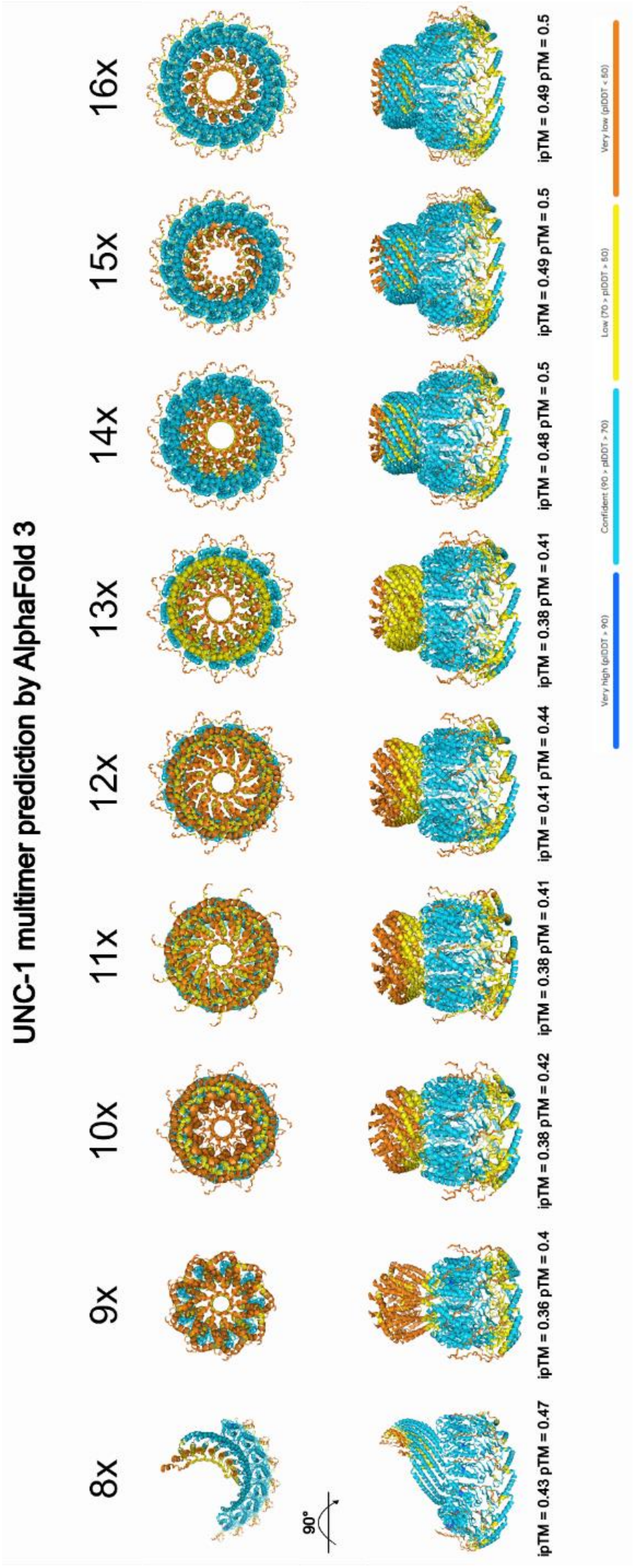


**Fig. S10. AF3 structural models of different UNC-1 multimers.** Different copy numbers of UNC-1, as indicated, were modelled by AF3. Any number above n=8 produced ring like assemblies. Confidence scores (ipTM - integrated predicted template modeling), and colour-coded plDDT (predicted local distance difference test, a per-residue confidence metric) values of the structural models are indicated.

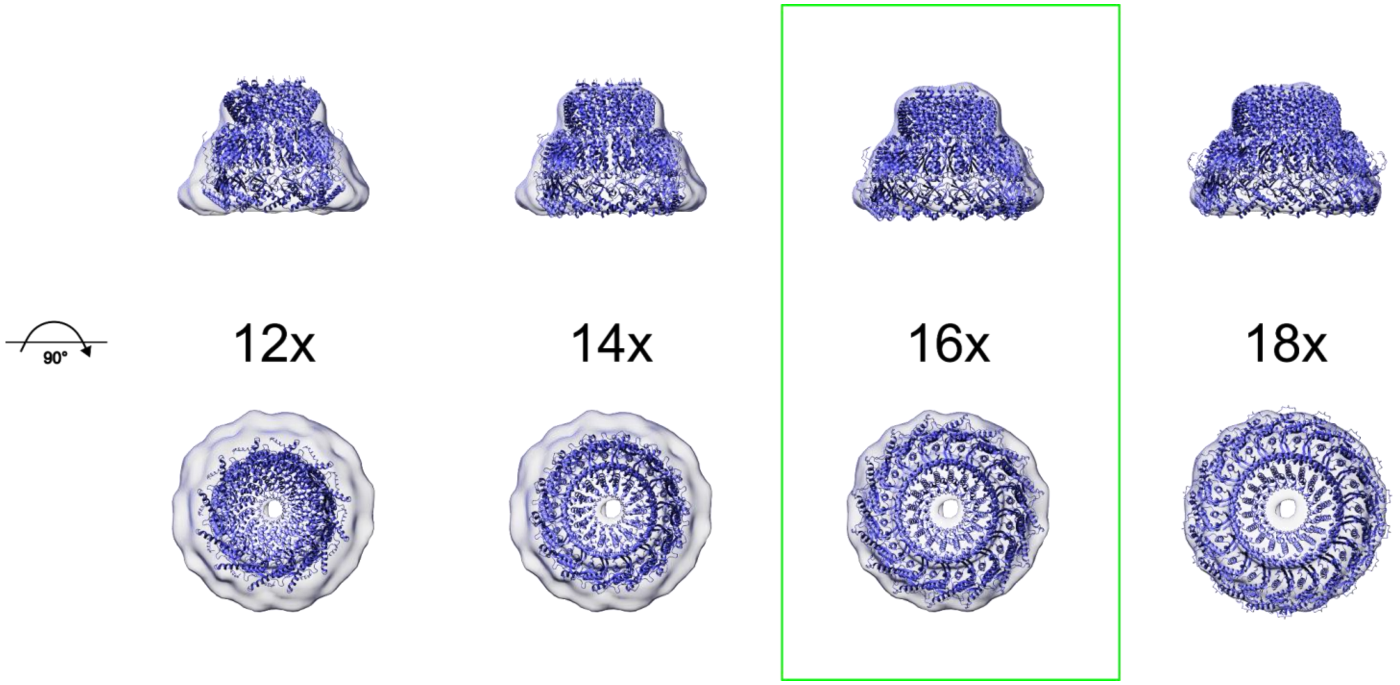


**Fig. S11. Fitting of different AF3 structural models of UNC-1 multimers into the *in situ* obtained cap surface map.** Different copy numbers of UNC-1, as indicated, were modelled by AF3, and fitted into the cap surface map. Side (upper row) and top (lower row) views are shown.

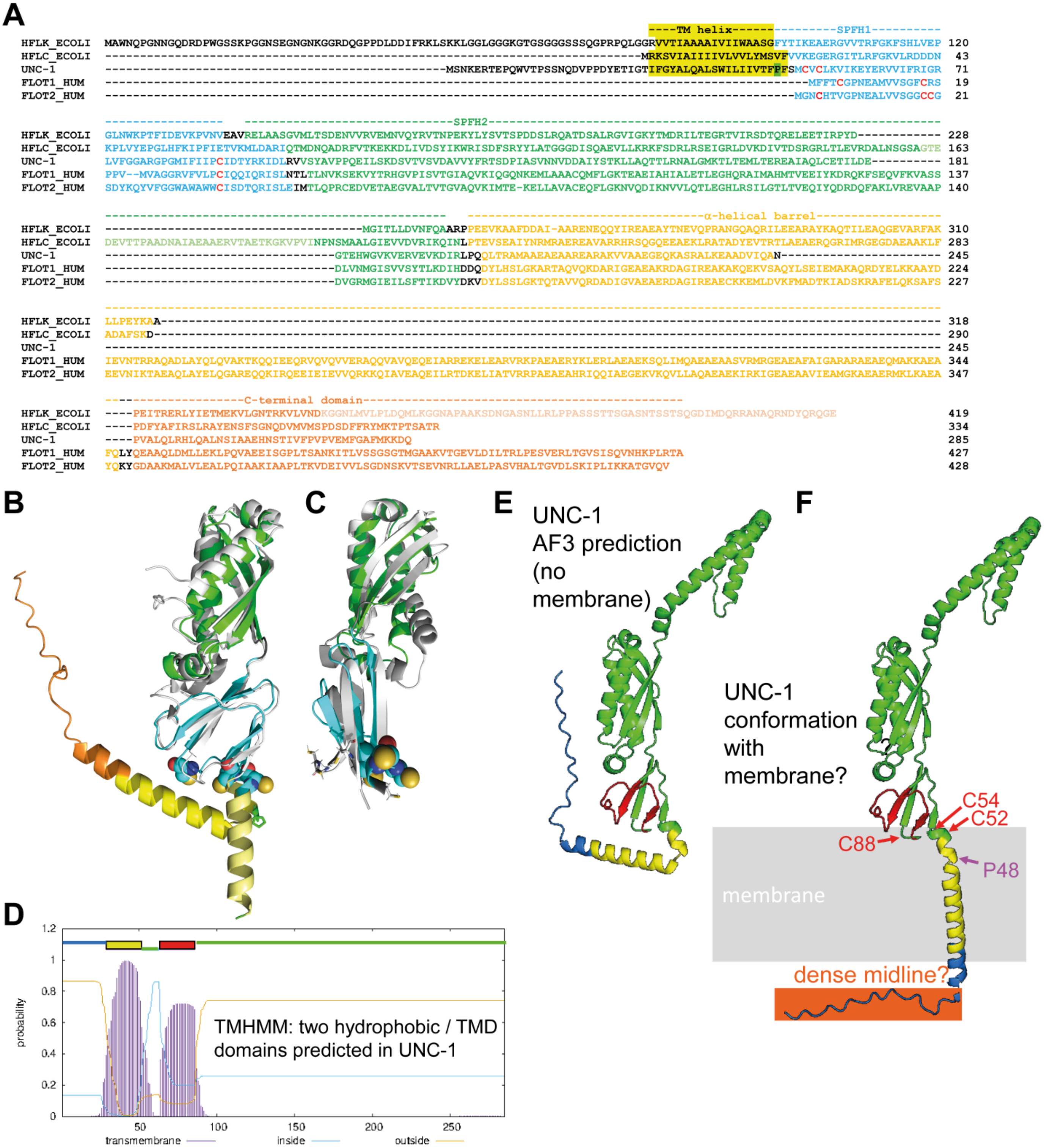


**Fig. S12. Alignment of UNC-1 with flotillins and HflC/K, and putative placement of TM domain of UNC-1. (A)** Alignment of UNC-1 with bacterial HflC/K and human Flotillins 1 and 2, based on structural features and the UNC-1 AF3 model. Pale colored sequence stretches are not visible in the biological structures. **(B)** Structural alignment of the N-terminal half of UNC-1 (domain colors as in the sequence alignment in A) and HflC (white). The TM helx of HflC is shown in pale yellow. The TM helix, predicted by TMHMM (D) is pointing upwards in the AF3 model. **(C)** Structural alignment of the N-terminal half of UNC-1 (domain colors as in the sequence alignment in A) and Flotilin2 (white). Cysteins that are palmytoylated in Flotillin are shown as sticks, equivalent cysteins in UNC-1 as spheres. **(D)** TMHMM predicts two hydrophobic / TM domains in UNC-1. **(E)** Predicted conformation of UNC-1 based on AF3 model. **(F)** Putative arrangement of the UNC-1

TM1 as traversing the membrane, with the N-terminal 29 amino acids possibly forming the dense midline seen in sub tomogram averages of capped GJs.

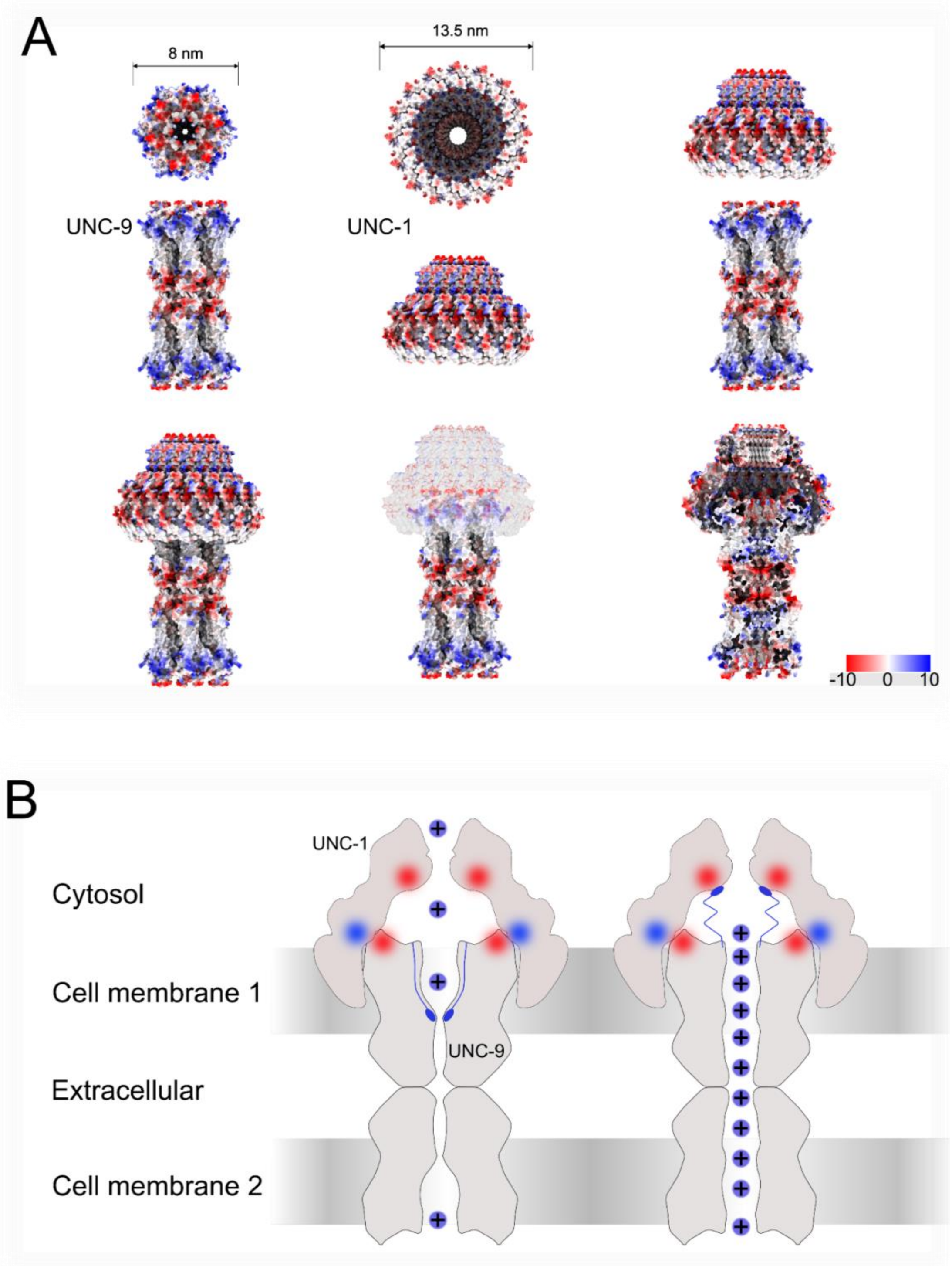


**Fig. S13. Electrostatic surface potentials and putative interactions of UNC-9 and UNC-1 multimeric models, as well as a model hypothesis for UNC-1 cap function. (A)** AF3 models of 12x UNC-9 and 16x UNC-1 were analyzed for their surface potentials, indicated in color score. Dimensions of the structures are indicated. Structures were automatically fitted in ChimeraX using the 3D rendering of the cap channel complex' surface map as a scaffold, highlighting possible interaction layers in the circular assemblies. **(B)** Model for how UNC-1 may function in the context of GJ gating. Color indicates surface potential, as derived from (A). The positively charged UNC-9 N-termini are located in the pore opening, thus closing the channels when there is a transjunctional potential difference. When the potential difference is close to zero, N-termini are released from their binding site in the pore and may then be arranged in the hollow interior of the UNC-1 cage, possibly binding to negative charges in the top part of the inside of the cap, as indicated.

**Table S1.**

**Statistics of open and closed GJ channels, analyzed in this work.** GJ channels were observed in three different tomograms. Shown are total numbers, as well as GJs in putative open or closed states, in top and side views, as indicated.

| | | Side Views | Top Views | Total channels |
|---|---|---|---|---|
| Open GJs | Tomogram 1 | 80 | 84 | 164 |
| | Tomogram 2 | 123 | 129 | 252 |
| | Tomogram 3 | 69 | 77 | 146 |
| | Total | 272 | 290 | 562 |
| Closed GJs | Tomogram 1 | 84 | 0 | 84 |
| | Tomogram 2 | 180 | 79 | 259 |
| | Tomogram 3 | 69 | 70 | 139 |
| | Total | 333 | 149 | 482 |

**Table S2.**

**Statistics of GJ exhibiting cytosolic 'cap' structures.** Capped GJs were observed in seven different tomograms. Shown are total numbers, as well as numbers of GJs with one or two caps. Also given are the ratios of single capped and double capped channels per tomogram, and overall.

| | Channels with one cap | Channels with two caps | Total channels | Ratio single capped channels | Ratio double capped channels |
|---|---|---|---|---|---|
| Tomogram 1 | 15 | 12 | 27 | 0.56 | 0.44 |
| Tomogram 2 | 74 | 31 | 105 | 0.7 | 0.3 |
| Tomogram 3 | 17 | 4 | 21 | 0.81 | 0.19 |
| Tomogram 4 | 6 | 4 | 10 | 0.6 | 0.4 |
| Tomogram 5 | 36 | 10 | 46 | 0.78 | 0.22 |
| Tomogram 6 | 23 | 0 | 23 | 1 | 0 |
| Tomogram 7 | 25 | 1 | 26 | 0.96 | 0.04 |
| Total | 196 | 62 | 258 | 0.76 | 0.24 |

**Movie S1.**

**Segmentation of cryo-ET slices, highlighting arrangement of cell membranes and organelles surrounding a gap junction.** Different compartements are color coded and highlighted in the movie. Zoom-in on the junction shows capped and uncapped channels next to each other, forming a gap junction assembly. Last, the movie shows a structural representation of the UNC-1 and UNC-9 multimeric models derived from AlphaFold 3 fitted into the cap junction.
